# A reactivated thalamocortical plasticity window promotes learning and is reshaped by experience

**DOI:** 10.64898/2026.08.27.747577

**Authors:** Hyesoo Jie, Nikorn Pothayee, Zhi-De Deng, Kathryn Sharer, Alan P. Koretsky

**Author notes:** Correspondence: Hyesoo Jie; Alan P. Koretsky.

## Abstract

Adult sensory loss can reactivate critical-period-like thalamocortical plasticity, but whether this reactivation defines a temporally gated circuit state that facilitates learning and is reciprocally shaped by experience remains unknown. Here we define its *in vivo* trajectory and functional consequences in adult mouse barrel cortex. Infraorbital nerve transection opened a transient window of enhanced layer 4 thalamocortical gain. Training during this window lowered whisker-detection thresholds and promoted learning by accelerating the transition to stable performance. Local GluN2B blockade prevented both cortical potentiation and the learning advantage, linking critical-period-associated plasticity mechanisms to adaptive behavior in the adult brain. Neuropixels recordings showed that weak inputs preferentially increased neuronal responses, whereas strong inputs produced sharper temporal coding. The relationship was reciprocal: experience reshaped the trajectory of this circuit state, with training before the normal peak advancing the emergence of potentiation, training during the active window prolonging the potentiated state, and training after closure failing to reinstate potentiation. State prolongation accompanied more persistent sensory memory. These findings establish a reciprocal, timing-dependent interaction between endogenous plasticity and experience, revealing a general principle by which adult circuits can convert transient plastic potential into adaptive behavioral change and informing strategies that align training with periods of heightened plasticity.

## Introduction

Adult sensory systems must preserve stable representations while retaining the capacity to adapt after input loss. This balance is central to both sensory function and recovery because deprivation can reorganize spared or cross-modal cortical circuits, with outcomes ranging from adaptive compensation to abnormal percepts and pain.^1–12^ A key question is therefore not simply whether adult circuits remain plastic, but whether plasticity is organized into temporally defined states with distinct consequences for perception and learning. Through this lens, recovery depends not merely on the magnitude of available plasticity, but on the precise temporal alignment between experience and a receptive circuit state capable of translating plastic potential into adaptive behavioral changes.

The thalamocortical input to layer 4 (L4) of primary sensory cortex provides a tractable entry point for testing this state-dependent view. In the whisker system, L4 barrel cortex is the principal cortical recipient of ventral posteromedial thalamic (VPM) input and links thalamic recruitment to tactile-guided behavior.^13–20^ Unilateral infraorbital nerve transection (IONX), which removes trigeminal input from one whisker pad, reactivates critical-period-like plasticity at spared adult thalamocortical (TC) synapses. Previous work established GluN2B-dependent, silent-synapse-like potentiation of spared TC inputs after IONX, its dependence on spared-whisker experience, and downstream changes across sensorimotor circuits.^21–23^ What remained unknown was whether this reactivated synaptic plasticity enhances sensory-guided learning and perception *in vivo*, how it reshapes the cortical representation of surviving sensory input, and whether experience can, in turn, alter the trajectory of this plastic state. Answering these questions would establish the functional significance of reactivated TC plasticity by linking its synaptic expression to cortical sensory coding and adaptive learning, while revealing how experience feeds back to reshape the trajectory of the plasticity window.

Here we align whisker-detection training to physiologically defined phases of post-denervation TC enhancement and combine longitudinally staged physiology, latent-state modeling of learning, local GluN2B blockade, and anatomically validated Neuropixels recordings. We then shift training across postoperative phases to test the converse question: whether experience merely exploits the reactivated window or reshapes its trajectory. We find that sensory loss opens a transient state that enhances weak-input recruitment, lowers perceptual threshold, and accelerates stabilization of learned performance. Experience, in turn, alters the trajectory of this state in a phase-dependent manner—advancing an emerging window, prolonging an active window, and failing to restore it after closure. Across molecular, circuit, and behavioral levels, these experiments identify a reciprocal timing principle: a transient plastic state changes what the adult circuit can learn, and appropriately timed learning feeds back to change the trajectory of that state. This framework unifies critical-period-like reactivation, sensory gain, learning stabilization, and memory persistence within a single state-dependent model of adult adaptation.

## Results

### Sensory loss reactivates a transient window of enhanced thalamocortical gain

We first determined whether post-denervation TC enhancement is sustained or instead occupies a discrete temporal window. Whisker-evoked L4 responses were mapped in independent sham and IONX cohorts at predefined postoperative intervals spanning the emergence, peak, persistence, and closure of enhancement (Fig. 1a). L4 local field potential (LFP) peak amplitude was used as an operational readout of TC gain. A fixed 6.7° whisker deflection—equivalent in the present geometry to the 470-µm displacement that previously produced robust L4 responses and clear IONX–sham separation^23^—provided a standardized physiological assay across time.

**Fig. 1.**
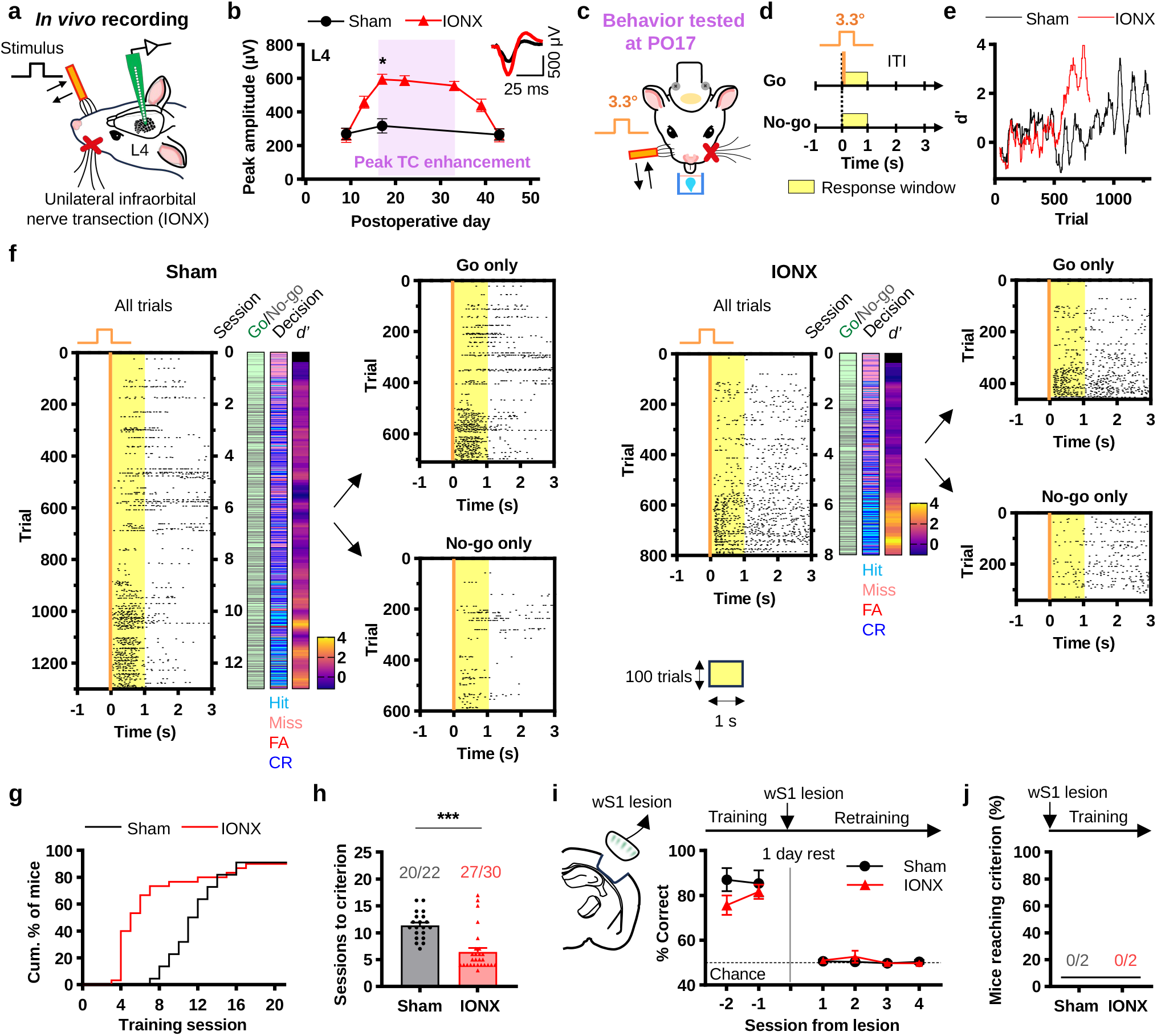
IONX reactivates a transient thalamocortical plasticity window that promotes sensory learning. (a) Schematic of IONX and whisker-evoked recording from L4 barrel cortex on the spared-whisker side. (b) Whisker-evoked L4 LFP peak amplitudes in independent postoperative cohorts. Sham: PO9, n = 6 mice; PO17 and PO43, n = 5 mice. IONX: PO9, PO13, PO17, PO22, PO39, and PO43, n = 6 mice; PO33, n = 4 mice. Time-matched Sham–IONX comparisons at PO9, PO17, and PO43 used two-sided Mann–Whitney U-tests with Holm correction; only PO17 differed (U = 0, raw P = 0.00433, adjusted P = 0.01299; PO9 and PO43, adjusted P = 1.0). Insets, representative L4 LFP traces; scale bars, 25 ms and 500 µV. (c) Primary behavioral training began at PO17, the peak of IONX-induced TC enhancement. (d) Head-fixed Go/No-go task. On Go trials, a 3.3° deflection was delivered, and licking within 1 s was rewarded; licking on unstimulated No-go trials was a false alarm. (e) Representative trial-resolved sensitivity (d′) trajectories during acquisition. (f) Representative lick rasters, classified as hits, misses, false alarms, or correct rejections. (g) Cumulative criterion attainment was earlier after IONX (Sham, n = 22 mice; IONX, n = 30 mice; log-rank Mantel–Cox: χ^2^ = 4.047, P = 0.0442; hazard ratio, 1.91; 95% confidence interval, 1.017–3.604). (h) Sessions to criterion among learners (Sham, 11.35 ± 0.56 sessions, n = 20 mice; IONX, 6.44 ± 0.76 sessions, n = 27 mice; two-sided Mann–Whitney U-test, U = 74.5, P = 2.34 × 10^−5^). (i) Post-learning wS1 lesion reduced retrained performance to chance (n = 3 mice/group). (j) Pre-learning wS1 lesion prevented acquisition (0/2 mice/group). Data are mean ± SEM. Points in h represent mice. *P < 0.05, **P < 0.01, ***P < 0.001. Abbreviations: IONX, infraorbital nerve transection; L4, layer 4; TC, thalamocortical; PO, postoperative day; LFP, local field potential; wS1, whisker primary somatosensory cortex.

L4 responses were stable across sampled time points in sham mice but increased transiently after IONX. Responses rose after denervation, peaked at postoperative day 17 (PO17), remained elevated through PO33, and returned to sham-like levels by PO43 (Fig. 1b). Time-matched comparisons detected no IONX–sham difference at PO9 or PO43, whereas L4 LFP peak amplitude was markedly greater at PO17 (sham, 317.62 ± 42.04, n = 5 mice; IONX, 594.81 ± 29.56, n = 6 mice; two-sided Mann–Whitney U-test, U = 0, P = 0.00433; Holm-adjusted P = 0.01299). Thus, sensory loss did not chronically elevate cortical responsiveness; it opened a transient window of enhanced TC drive that peaked at PO17 and closed by PO43. This temporally bounded profile establishes post-denervation plasticity as a discrete circuit state with a defined lifespan, allowing us to directly test whether behavioral benefit depends on the timing of experience relative to the underlying plastic window.

This physiological trajectory defined the experimental time axis for the behavioral studies. We used PO17 for primary training, then independently shifted training onset to earlier, active, and post-closure phases to test whether experience could advance, prolong, or reinstate the TC state (Fig. 1c and Supplementary Fig. 1a,b).

### A reactivated plasticity window promotes sensory learning

To test whether the active TC state changes learning capacity, mice began a head-fixed Go/No-go whisker-detection task at PO17. A custom coil-driven actuator delivered calibrated 2-ms deflections, and mice were trained to respond to detection by licking within a 1-s response window (Fig. 1c,d, Supplementary Fig. 2a–d, and Supplementary Movie 1). Behavioral training used a 3.3° deflection, whereas the physiological time-course assay used 6.7°. The 3.3° stimulus lies within the dynamic, non-ceiling portion of the input–output relationship^23^ and matches the readily detectable starting range of an established psychometric paradigm.^24^ Learning criterion was ≥80% hit rate and ≤30% false-alarm rate for two consecutive sessions, with training limited to 21 sessions.

IONX mice acquired the 3.3° task earlier than sham controls. Trial-resolved behavior showed a more rapid rise in stimulus-locked licking, and cumulative criterion attainment shifted toward earlier sessions (Fig. 1e–g). With non-learners treated as right-censored observations, median time to criterion was 11.5 sessions in sham mice and 5.0 sessions after IONX (log-rank Mantel–Cox test, χ^2^ = 4.047, d.f. = 1, P = 0.0442; hazard ratio, 1.91; 95% confidence interval, 1.017–3.604; Fig. 1g). Among learners, IONX mice likewise required fewer sessions (sham, 11.35 ± 0.56 sessions, n = 20 mice; IONX, 6.44 ± 0.76 sessions, n = 27 mice; two-sided Mann–Whitney U-test, U = 74.5, P = 2.34 × 10^−5^; Fig. 1h).

Given evidence that barrel cortex contributes to whisker perception but is not uniformly required across tactile tasks,^20,25^ we next asked whether the IONX-associated learning advantage required the spared whisker primary somatosensory cortex (wS1) pathway rather than alternative cues or nonspecific response strategies. Lesioning contralateral wS1 after learning reduced performance to chance in both groups (n = 3 mice per group; Fig. 1i and Supplementary Fig. 3d,e), whereas lesions before PO17 training prevented criterion attainment (0/2 mice per group; Fig. 1j and Supplementary Fig. 3b,c). Removing the trained whiskers after learning likewise reduced performance to chance (Supplementary Fig. 3f,g). In paired VPM–L4 recordings, IONX left thalamic responses unchanged but increased cortical responses and produced a 1.7-fold increase in the VPM-to-L4 amplification slope (sham, 2.12 ± 0.48; IONX, 3.61 ± 0.36; Supplementary Fig. 4). Together, these controls localize the behavioral advantage to cortical processing of spared tactile input and support enhanced transformation of intact thalamic signals rather than increased VPM drive. Thus, pathway reactivation enhances cortical processing of surviving sensory input, providing a circuit substrate for adaptive compensation after unilateral sensory loss.

### Reactivated plasticity promotes learning by accelerating stabilization

Faster criterion attainment could reflect earlier learning onset, faster stabilization after learning begins, or changes in licking-related motor output. Early nonspecific licking was unlikely to explain the group difference: first-day lick probability accounted for only 3% of the variance in sessions to criterion (R^2^ = 0.030, P = 0.0240). Excluding the small number of high-licking animals did not remove the IONX-associated advantage (Supplementary Fig. 5). We therefore applied a three-state hidden Markov model (HMM), adapted from latent-state analyses of behavioral learning,^26–29^ to infer pre-learning, active-learning, and post-learning states from block-wise hit and false-alarm patterns (Fig. 2a,c).

**Fig. 2.**
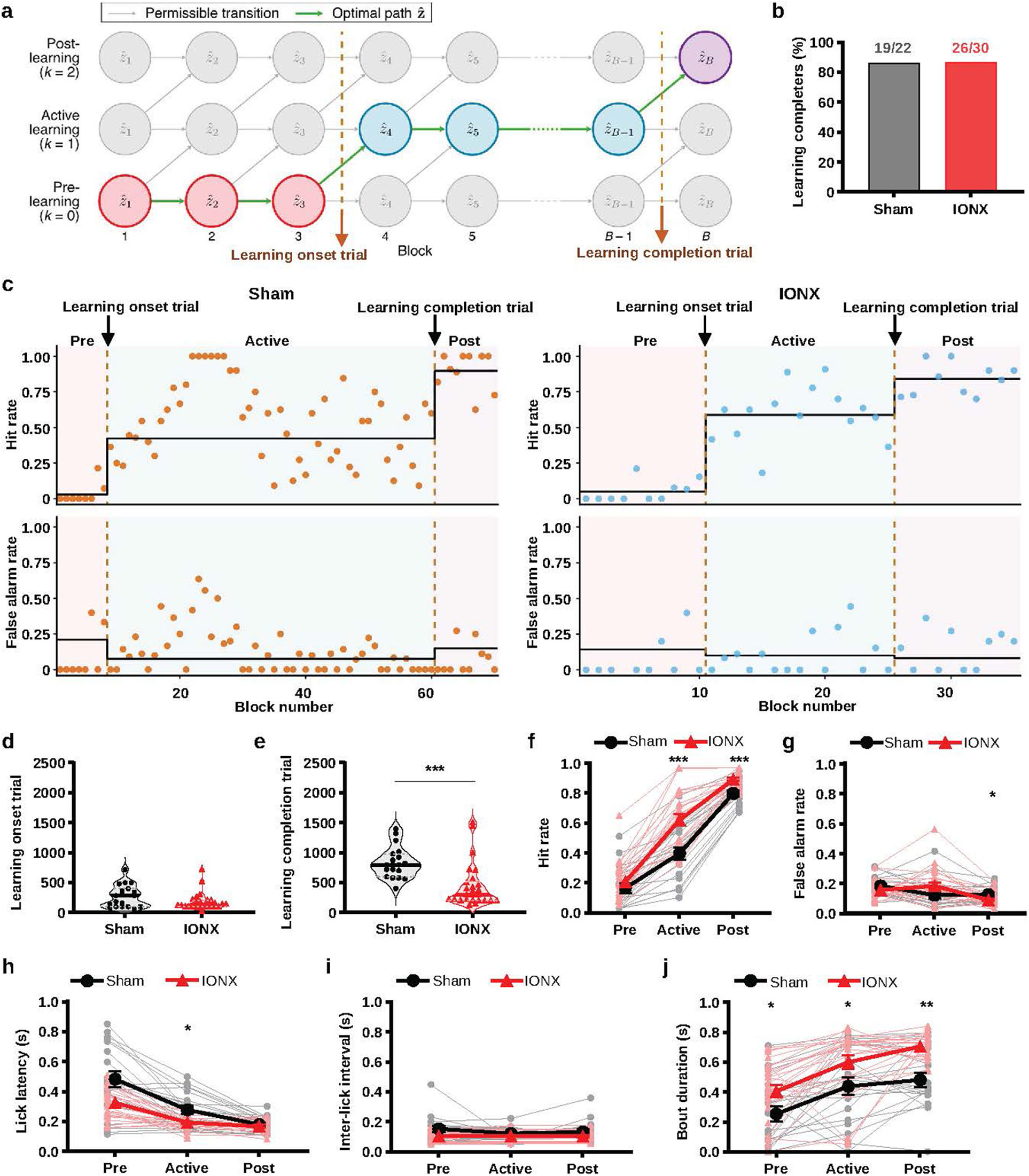
Reactivated plasticity promotes learning by accelerating stabilization of learned performance. (a) Three-state HMM schematic for pre-learning, active-learning, and post-learning states inferred from 20-trial blocks. Gray arrows show permitted transitions, the green path shows the Viterbi-decoded sequence, and dashed lines mark learning onset and completion. (b) HMM-defined completers (Sham, 19/22 mice; IONX, 26/30 mice; two-sided Fisher’s exact test, P = 1.0). (c) Representative HMM fits with block-wise hit and false-alarm rates and fitted state probabilities. (d) Learning onset (Sham, 272.63 ± 42.45 trials, n = 19 mice; IONX, 177.69 ± 28.72 trials, n = 26 mice; two-sided Mann–Whitney U-test, U = 316, P = 0.1108). (e) HMM-defined learning completion (Sham, 831.58 ± 60.47 trials, n = 19 mice; IONX, 440.77 ± 71.85 trials, n = 26 mice; two-sided Mann–Whitney U-test, U = 423.5, P = 5.08 × 10^−5^). (f) HMM-estimated state-specific hit probabilities were higher after IONX during active learning and post-learning (Holm-adjusted P = 0.00055 and 0.00085), but not pre-learning. (g) HMM-estimated state-specific false-alarm probabilities differed only during post-learning (Holm-adjusted P = 0.03218). (h) Hit-trial lick latency (Sham, n = 19/19/19 mice; IONX, n = 26/26/23 mice) was shorter after IONX during active learning (Holm-adjusted P = 0.02173). (i) Inter-lick interval (Sham, n = 19/19/19 mice; IONX, n = 25/26/23 mice); no within-state group comparison survived Holm correction. (j) Bout duration (Sham, n = 19/19/19 mice; IONX, n = 26/26/23 mice) was longer after IONX across states (Holm-adjusted P = 0.02536, 0.02536, and 0.00206). For h–j, n values are listed in pre-/active-/post-learning order. Data are mean ± SEM. Stars in f–j denote prespecified within-state group contrasts from mixed-effects models, Holm-corrected within outcome. *P < 0.05, **P < 0.01, ***P < 0.001. Abbreviations: HMM, hidden Markov model; IONX, infraorbital nerve transection.

The proportion of HMM-defined completers did not differ between groups (sham, 19/22; IONX, 26/30; two-sided Fisher’s exact test, P = 1.0; Fig. 2b). Learning onset was also not significantly shifted (sham, 272.63 ± 42.45 trials, n = 19 mice; IONX, 177.69 ± 28.72 trials, n = 26 mice; two-sided Mann–Whitney U-test, U = 316, P = 0.1108; Fig. 2d). In contrast, HMM-defined learning completion occurred substantially earlier after IONX (sham, 831.58 ± 60.47 trials; IONX, 440.77 ± 71.85 trials; U = 423.5, P = 5.08 × 10^−5^; Fig. 2e). The principal behavioral effect was therefore a compressed transition from active learning to stable performance, rather than a detectable advance in learning onset. This distinction identifies stabilization of learned behavior as the component of acquisition most strongly facilitated by the reactivated state.

State-resolved response probabilities supported this interpretation. Fitted hit probability was higher after IONX during active learning and post-learning, but not pre-learning (Holm-adjusted P = 0.00055, 0.00085, and 0.09403, respectively; Fig. 2f). False-alarm probability followed different state trajectories in the two groups and differed significantly only after learning (Holm-adjusted P = 0.03218; Fig. 2g). Thus, earlier stabilization was associated primarily with stronger stimulus-evoked responding rather than a generalized tendency to lick.

Lick-output dynamics changed in parallel but could be separated from the stabilization effect. Hit-trial latency was similar before learning, transiently shorter after IONX during active learning, and converged after learning (Holm-adjusted P = 0.02173 during active learning; Fig. 2h). Inter-lick interval showed an overall group effect, but no state-wise contrast survived Holm correction (Fig. 2i), whereas lick bouts were longer after IONX across states (Holm-adjusted P = 0.02536, 0.02536, and 0.00206; Fig. 2j). These motor-output differences therefore accompany, but do not account for, the main learning phenotype: earlier stabilization of a sensory-guided stimulus–action association. The reactivated TC state therefore changes the efficiency with which sensory evidence is converted into stable learned behavior, rather than simply increasing response vigor.

### Barrel-cortex GluN2B signaling links potentiation to the learning advantage

Previous studies established that IONX restores GluN2B-dependent, silent-synapse-like potentiation at spared adult TC synapses.^21,22^ We asked whether local GluN2B signaling in barrel cortex is required *in vivo* both for the potentiated L4 state and for its behavioral advantage. The selective GluN2B antagonist Ro25-6981 or vehicle was infused into wS1 during PO0–PO14; whisker-detection training began at PO17, followed by L4 physiology after learning (Fig. 3a,b,d).

**Fig. 3.**
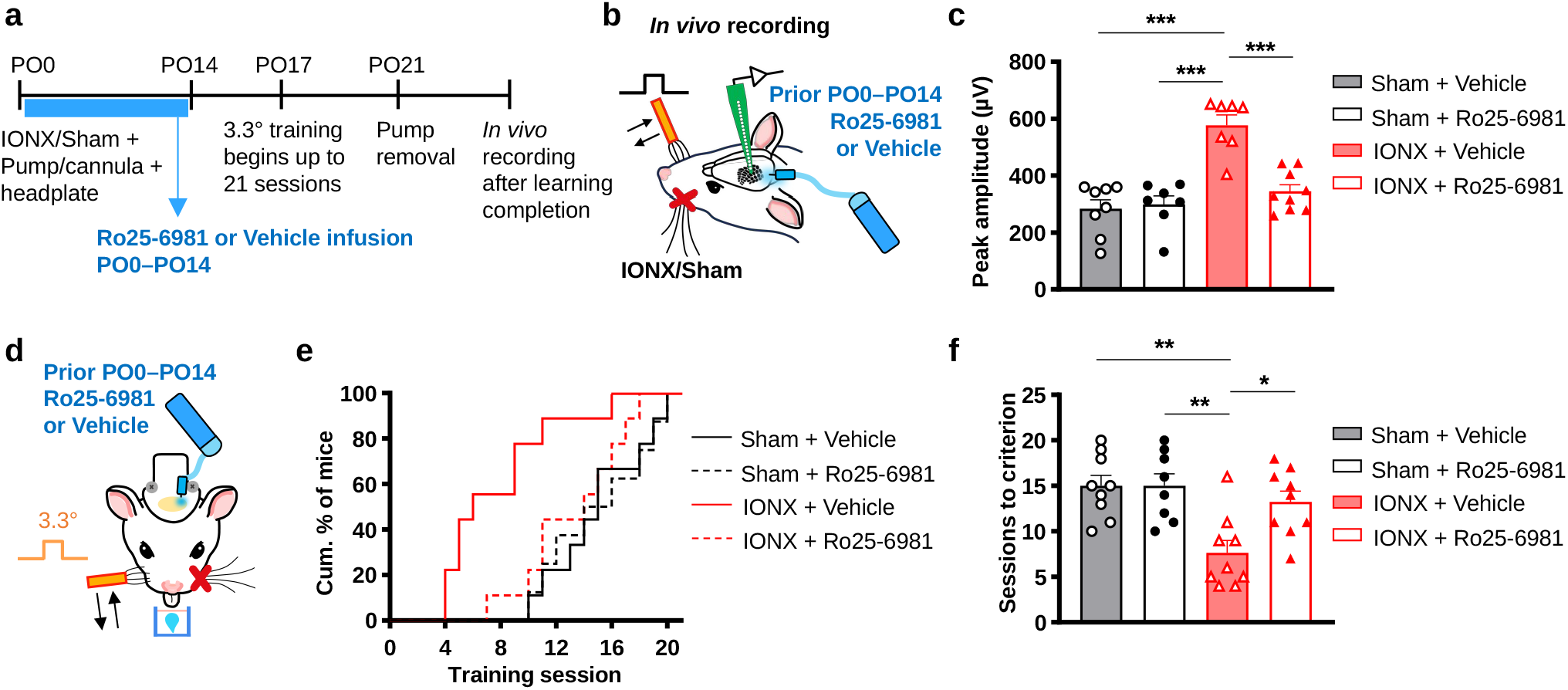
Local GluN2B signaling links reactivated thalamocortical potentiation to the learning advantage. (a) Local Ro25-6981 or vehicle infusion during the early postoperative induction window after IONX or sham surgery. (b) *In vivo* whisker-evoked L4 recording after learning completion. (c) L4 TC peak amplitude (Sham + Vehicle, n = 8 mice; Sham + Ro25-6981, n = 7 mice; IONX + Vehicle, n = 7 mice; IONX + Ro25-6981, n = 9 mice). Ro25-6981 suppressed IONX-induced enhancement (two-way ANOVA, surgery × drug: F(1,27) = 16.78, P = 3.43 × 10^−4^; Tukey-adjusted comparisons of IONX + Vehicle with each other group, all P < 0.0001). (d) Behavioral testing after PO0–PO14 infusion. (e) Cumulative criterion attainment differed across groups (Sham + Vehicle, n = 9 mice; Sham + Ro25-6981, n = 8 mice; IONX + Vehicle, n = 9 mice; IONX + Ro25-6981, n = 9 mice; log-rank Mantel–Cox, χ^2^ = 17.51, P = 5.56 × 10^−4^). Ro25-6981 delayed learning after IONX (IONX + Vehicle versus IONX + Ro25-6981, P = 0.0166). (f) Sessions to criterion differed across groups (Sham + Vehicle, n = 9 mice; Sham + Ro25-6981, n = 8 mice; IONX + Vehicle, n = 9 mice; IONX + Ro25-6981, n = 9 mice; one-way ANOVA, F(3,31) = 7.63, P = 5.82 × 10^−4^; Tukey-adjusted comparisons of IONX + Vehicle with Sham + Vehicle, Sham + Ro25-6981, and IONX + Ro25-6981: P = 0.0013, 0.0018, and 0.0179). Data are mean ± SEM. Points in c and f represent mice. *P < 0.05, **P < 0.01, ***P < 0.001. Abbreviations: IONX, infraorbital nerve transection; TC, thalamocortical; L4, layer 4; GluN2B, N-methyl-D-aspartate receptor subunit 2B.

IONX increased whisker-evoked L4 responses in vehicle-treated mice, whereas Ro25-6981 reduced this enhancement to sham-like levels (two-way ANOVA, surgery × drug interaction: F(1,27) = 16.78, P = 3.43 × 10^−4^; IONX + vehicle versus sham + vehicle, adjusted P < 0.0001; IONX + vehicle versus IONX + Ro25-6981, adjusted P < 0.0001; Fig. 3c). Ro25-6981 had little effect in sham mice, arguing against generalized suppression of baseline cortical transmission. Independent dye-infusion and histological validation confirmed overlap with the intended wS1 target and cannula placement in all 35 pharmacology animals (Supplementary Fig. 6).

Ro25-6981 also attenuated the IONX-associated learning advantage. Criterion-attainment curves differed across the four treatment groups (overall log-rank Mantel–Cox test, χ^2^ = 17.51, d.f. = 3, P = 5.56 × 10^−4^; Fig. 3e): vehicle-treated IONX mice learned earlier than vehicle-treated sham mice, whereas Ro25-6981 delayed criterion attainment after IONX (IONX + vehicle versus sham + vehicle, P = 0.00358; IONX + vehicle versus IONX + Ro25-6981, P = 0.0166). Sessions to criterion showed the same pattern (one-way ANOVA, F(3,31) = 7.63, P = 5.82 × 10^−4^; Tukey-adjusted P = 0.0013 and 0.0179 for the corresponding comparisons; Fig. 3f). Local GluN2B signaling during the induction period is therefore required for both later L4 potentiation and accelerated learning, directly linking the reactivated TC state to behavioral benefit. Thus, re-engagement of a critical-period-associated molecular mechanism is not merely a physiological signature of deafferentation; it determines whether the adult circuit expresses the learning advantage associated with the reactivated state.

### Sensory loss enhances weak-input recruitment, lowers perceptual threshold, and sharpens strong-input timing

We next asked how the reactivated state changes sensory processing across input intensities. Neuropixels recording sites in the whisker TC pathway were independently constrained by VPM barreloid anatomy, VPM axon labeling, tangential barrel mapping, probe tracks labeled with the lipophilic carbocyanine dye DiI, and current-source-density localization of the earliest L4 input sink (Fig. 4 and Supplementary Fig. 4a). Spike-count analyses used validated L4 single- and multi-unit activity sites, whereas temporal analyses used validated L4 units.

**Fig. 4.**
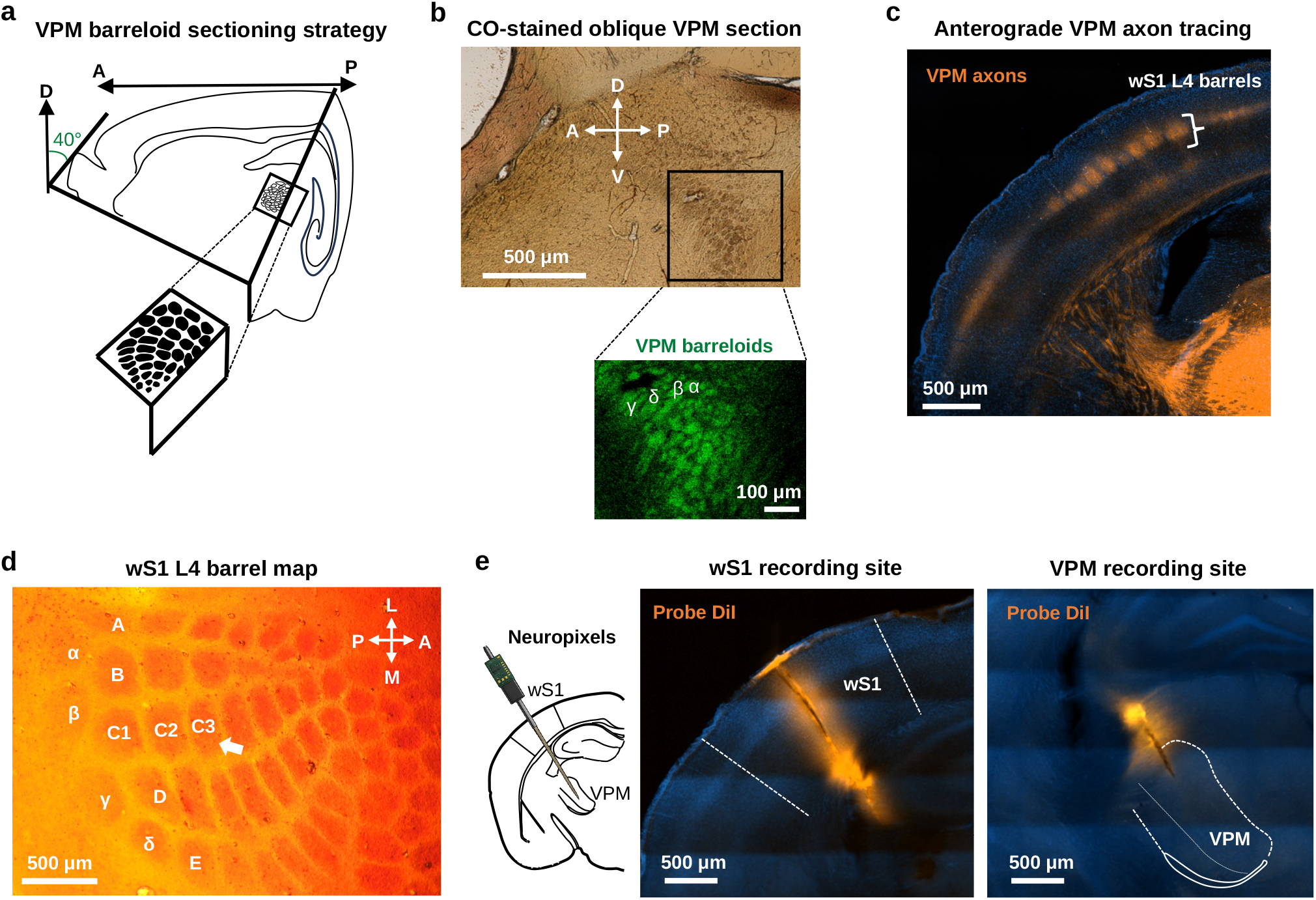
Anatomical validation of Neuropixels recording sites in wS1 and VPM. (a) Schematic of the ∼40° oblique sectioning plane used to visualize the VPM barreloid array, which is not readily resolved in standard coronal sections. The sectioning orientation was based on prior anatomical descriptions.^51^ (b) Representative cytochrome oxidase-stained oblique section obtained using the strategy in (a), showing VPM, with a higher-magnification green-autofluorescence view of the barreloid array. (c) Anterograde labeling of VPM axons in wS1 delineated the L4 barrel field and cortical recording zone. (d) Tangential cytochrome oxidase-stained section through wS1 L4 showing the barrel map and the associated probe track. (e) Representative DiI-labeled Neuropixels probe trajectories through wS1 and VPM. Scale bars: b, 500 µm (overview) and 100 µm (inset); c, 500 µm; d, 500 µm; e, 500 µm. Abbreviations: wS1, whisker primary somatosensory cortex; VPM, ventral posteromedial thalamic nucleus; L4, layer 4; DiI, lipophilic carbocyanine dye used for probe-track labeling.

If enhanced TC gain improves the representation of marginal sensory evidence, its behavioral effect should be largest for weak inputs. Consistent with this hypothesis, fixed-intensity training at PO17 revealed the strongest advantage at 1.8°: 2 of 11 sham mice reached criterion compared with 11 of 12 IONX mice (two-sided Fisher’s exact test, P = 0.000644; Fig. 5a,b). Time to criterion was likewise shorter after IONX at 1.8° (log-rank Mantel–Cox test, χ^2^ = 17.15, P = 3.45 × 10^−5^) and 3.3° (χ^2^ = 4.047, P = 0.0442), but not at the strong 8.3° stimulus (χ^2^ = 0.258, P = 0.612; Fig. 5c). The learning advantage was therefore greatest when sensory input was limiting, indicating that the reactivated state preferentially expands behavioral access to weak sensory evidence.

**Fig. 5.**
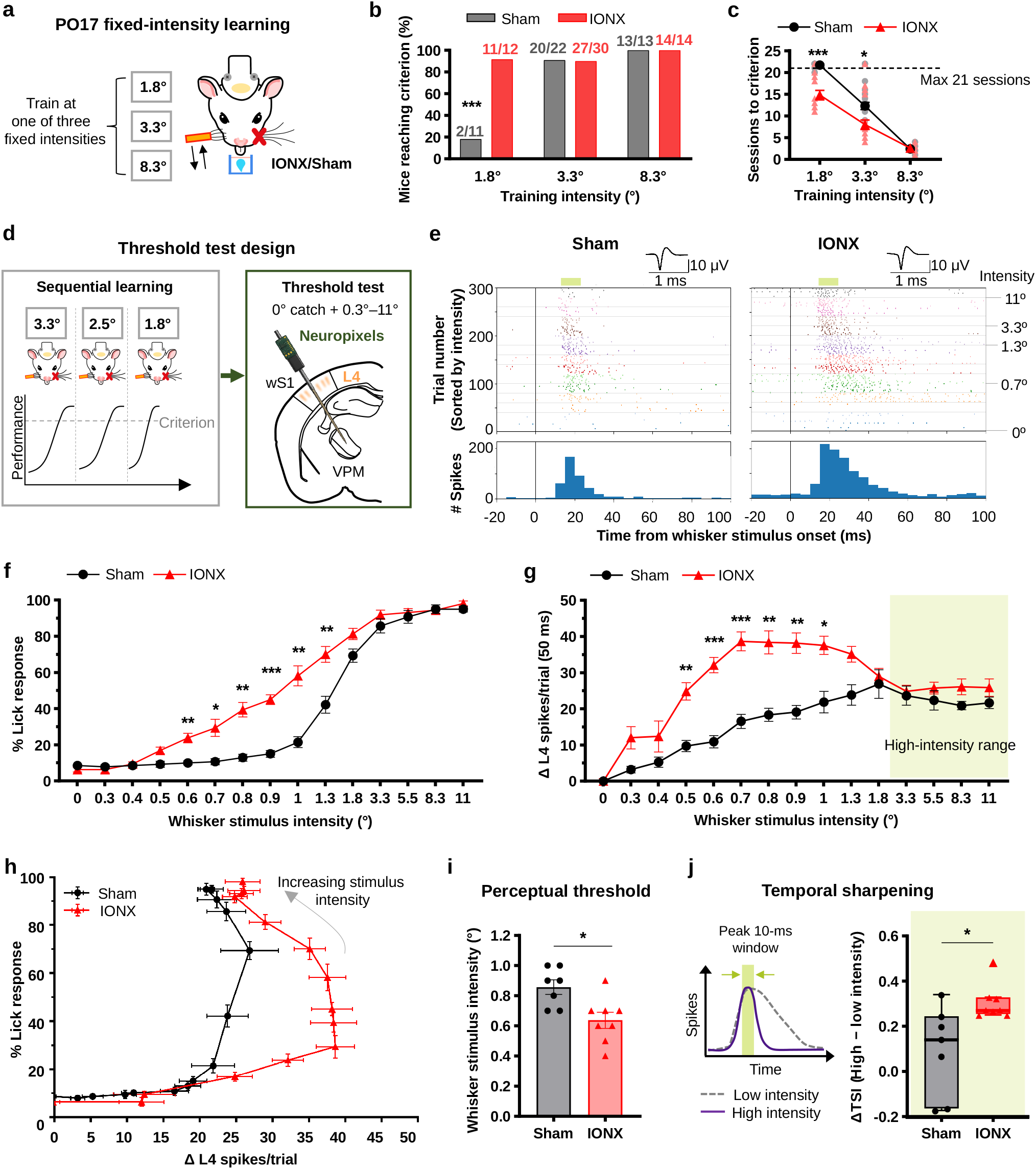
Reactivated thalamocortical plasticity enhances weak-input recruitment, lowers perceptual threshold, and sharpens strong-input timing. (a) PO17 fixed-intensity training (1.8°, 3.3°, or 8.3°). (b) Criterion attainment (Sham versus IONX): 2/11 versus 11/12 at 1.8°, 20/22 versus 27/30 at 3.3°, and 13/13 versus 14/14 at 8.3°; two-sided Fisher’s exact test at 1.8°, P = 0.000644. (c) Sessions to criterion; non-learners were right-censored at session 21 and plotted at 22. Log-rank tests: 1.8°, χ^2^ = 17.15, P = 3.45 × 10^−5^; 3.3°, χ^2^ = 4.047, P = 0.0442; 8.3°, χ^2^ = 0.258, P = 0.612. (d) Sequential 3.3°/2.5°/1.8° training preceded Neuropixels testing with catch trials and 14 deflections (0.3°–11°; 20 trials/condition). (e) Representative stimulus-aligned L4 spiking and unit waveforms. Light-green bars illustrate representative peak 10-ms windows, determined separately for each unit and intensity. Scale bars, 1 ms and 10 µV. (f) Psychometric performance (Sham, n = 7 mice; IONX, n = 8 mice). (g) Mouse-level L4 recruitment (Δ spikes/trial, 0–50 ms). Light-green shading denotes the high-intensity range used in j. Mixed ANOVAs for f and g showed significant group, intensity, and interaction effects (all P < 0.001). (h) Neural–behavioral trajectories. (i) Perceptual threshold was lower after IONX (Sham, 0.857 ± 0.048°; IONX, 0.638 ± 0.053°; n = 7 and 8 mice; two-sided exact permutation test, P = 0.0121). (j) Temporal sharpening. Left, TSI schematic showing the peak 10-ms response window. Right, ΔTSI (high minus low intensity) was greater after IONX (Sham, 0.094 ± 0.075; IONX, 0.306 ± 0.027; n = 7 and 8 mice; two-sided Mann–Whitney U-test, U = 7.0, P = 0.01399). Data are mean ± SEM; points represent mice. g and j show mouse averages across L4 activity sites and units, respectively. Stars in f–g denote Holm-adjusted intensity-wise comparisons; stars in i–j denote the tests reported above. *P < 0.05, **P < 0.01, ***P < 0.001. Abbreviations: IONX, infraorbital nerve transection; L4, layer 4; TSI, temporal sharpening index.

After sequential training at 3.3°, 2.5°, and 1.8°, mice were tested during Neuropixels recording with catch trials and 14 randomly interleaved whisker deflections from 0.3° to 11° (Fig. 5d). IONX shifted the psychometric function leftward (Fig. 5f), with significant effects of group, intensity, and their interaction (two-way mixed ANOVA; group, F(1,13) = 32.99, P = 6.78 × 10^−5^; intensity, F(14,182) = 399.53, P = 1.88 × 10^−128^; interaction, F(14,182) = 13.37, P = 1.98 × 10^−21^). *Post hoc* group differences were concentrated at 0.6°–1.3°.

The perceptual shift coincided with stronger L4 population recruitment at weak-to-intermediate intensities (Fig. 5e,g). Mouse-level spiking showed significant effects of group, intensity, and their interaction (group, F(1,13) = 40.63, P = 2.44 × 10^−5^; intensity, F(14,182) = 37.11, P = 7.88 × 10^−46^; interaction, F(14,182) = 6.63, P = 8.74 × 10^−11^), with greater IONX recruitment from 0.5° to 1.0°. Thus, the stimulus range showing the largest gain in cortical recruitment overlapped the range showing enhanced behavioral sensitivity.

To relate these measures within animals, we fit each intensity-ordered neural–behavioral trajectory with segmented total least-squares regression and defined the transition into the steep behavioral-response segment as an operational perceptual threshold (Fig. 5h and Supplementary Fig. 7a). Thresholds were lower after IONX (sham, 0.857 ± 0.048°, n = 7 mice; IONX, 0.638 ± 0.053°, n = 8 mice; two-sided exact permutation test based on the Mann–Whitney U statistic, U = 49.0, P = 0.0121; Fig. 5i), and a complementary manually assigned transition yielded concordant estimates (Supplementary Fig. 7b,c). Together, these findings indicate that the reactivated state preferentially enhances cortical recruitment and perceptual access when sensory input is weak.

We next asked whether, at stronger intensities, the reactivated state affected a complementary feature of sensory coding: the temporal organization of evoked spiking. To quantify this, we defined a temporal sharpening index (TSI) as the fraction of sensory-evoked spikes contained within the peak 10-ms window of the 0–100-ms post-stimulus response (Fig. 5e,j), motivated by established contributions of spike timing to sensory coding.^30,31^ For each mouse, ΔTSI was calculated as mean TSI at high intensities (3.3°, 5.5°, 8.3°, and 11°) minus mean TSI at low intensities (0.3°, 0.4°, and 0.5°). ΔTSI was greater after IONX (sham, 0.094 ± 0.075, n = 7 mice; IONX, 0.306 ± 0.027, n = 8 mice; two-sided Mann–Whitney U-test, U = 7.0, P = 0.01399; Fig. 5j and Supplementary Fig. 7d,e). Together with the weak-input recruitment effect, these recordings show that the reactivated state does not simply amplify cortical activity uniformly: it changes the representation of sensory evidence in an input-dependent manner, increasing recruitment when signals are weak and concentrating response timing when signals are strong.

### Experience reshapes the plasticity window in a phase-dependent manner

Finally, we asked whether learning simply benefits from the post-denervation state or also feeds back to control its time course. Independent sham and IONX cohorts began 3.3° training at PO4, PO10, PO17, or PO43 (Supplementary Fig. 1a), thereby sampling periods before the normal rise, near onset, at peak, and after closure of TC enhancement.

Training before the normal peak advanced the state. With PO4 onset, IONX mice reached criterion earlier than sham controls (P = 0.00288; Fig. 6a), and the physiological trajectory shifted forward: PO4-trained IONX mice already showed elevated L4 responses by PO13 relative to untrained IONX mice sampled at the same time (untrained, 458.28 ± 35.90; trained from PO4, 589.92 ± 27.48; n = 6 mice per group; two-sided Mann–Whitney U-test, U = 34, P = 0.00866; Fig. 6b). HMM-defined learning completion was also earlier after IONX in the PO4 cohort (P = 0.0132; Fig. 6c). Experience delivered during the emerging phase therefore advanced both behavioral stabilization and the expression of enhanced TC drive, showing that training can accelerate the emergence of endogenous plasticity when delivered before its normal peak.

**Fig. 6.**
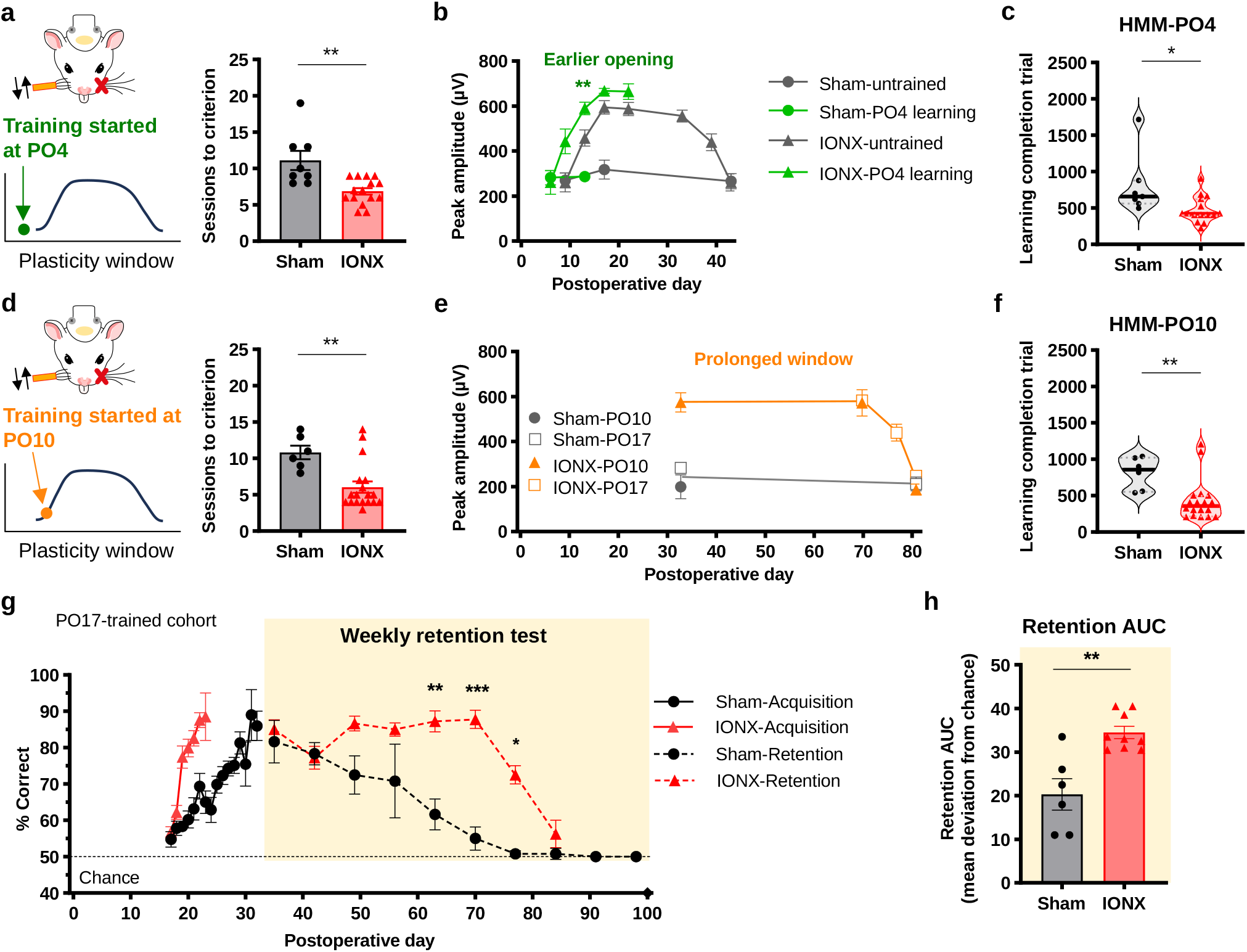
Experience reshapes the reactivated TC plasticity window in a phase-dependent manner. (a) PO4 acquisition was faster after IONX (Sham, n = 8 mice; IONX, n = 15 mice; U = 106, P = 0.00288). (b) PO4 training advanced L4 TC enhancement. Unless noted, n = 6 mice/cohort/time point; exceptions: untrained Sham PO17/PO43, n = 5 each; untrained IONX PO33, n = 4. PO13 trained versus untrained IONX: U = 34, P = 0.00866. (c) HMM completion after PO4 training (Sham, n = 7 mice; IONX, n = 15 mice; U = 88, P = 0.0132). (d) PO10 acquisition was faster after IONX (Sham, n = 6 mice; IONX, n = 18 mice; U = 94.5, P = 0.00676). (e) PO10 or PO17 training prolonged L4 TC enhancement (n = 6 mice/group/cohort; pooled n = 12 mice/group). (f) HMM completion after PO10 training (Sham, n = 6 mice; IONX, n = 18 mice; U = 96, P = 0.00541). (g) PO17 acquisition (solid) and retention (dashed). PO35–84 GEE: group at PO59.5, Wald χ^2^(1) = 31.21, P = 2.32 × 10^−8^; IONX slope, Wald χ^2^(1) = 9.79, P = 0.00176; interaction, Wald χ^2^(1) = 24.83, P = 6.27 × 10^−7^. Sham, n = 6 mice; IONX, n = 9 mice through PO70 and n = 4 mice at PO77/84 after planned terminal physiology. Stars, Holm-adjusted Welch’s t-tests. (h) Chance-referenced, time-normalized retention AUC (PO35–70) was greater after IONX (Sham, 20.33 ± 3.60, n = 6 mice; IONX, 34.50 ± 1.39, n = 9 mice; two-sided exact permutation test based on the Mann–Whitney U statistic, U = 5.5, P = 0.00879). Data are mean ± SEM; points are mice. a, c, d, and f: two-sided Mann–Whitney U-tests. *P < 0.05, **P < 0.01, ***P < 0.001. Abbreviations: IONX, infraorbital nerve transection; L4, layer 4; TC, thalamocortical; HMM, hidden Markov model; GEE, generalized estimating equation; AUC, area under the curve; PO, postoperative day.

Training near or within the active window instead produced prolonged enhancement. IONX mice trained from PO10 reached criterion earlier than sham controls (P = 0.00676; Fig. 6d), and training from PO10 or PO17 maintained L4 responses above sham levels beyond the time at which the untrained state had normally subsided (Fig. 6e). This persistence required sensory–reward contingency: when whisker stimulation and water reward were delivered without temporal pairing, PO70 responses returned to sham-like levels (Supplementary Fig. 8a,c,d). PO10-trained IONX mice also completed the HMM-defined transition earlier than sham controls (P = 0.00541; Fig. 6f). Thus, associative training can extend the physiological expression of an active TC state, converting a transient period of heightened gain into a more persistent circuit condition.

The influence of experience nevertheless had a temporal boundary. By PO43, untrained IONX responses had returned to sham-like levels, and training initiated at PO43 failed to reinstate L4 potentiation (Supplementary Fig. 8a,b). The same behavioral experience could therefore advance an emerging state or prolong an active state, but could not reopen the state once it had closed. This state dependence makes timing a functional variable: identical training produces different circuit outcomes depending on where it falls along the endogenous plasticity trajectory.

Prolongation of the learning-associated TC state was accompanied by more persistent sensory memory. In mice trained from PO17, retention trajectories differed across PO35–PO84 (Gaussian generalized estimating equation model; group at centered PO59.5, Wald χ^2^(1) = 31.21, P = 2.32 × 10^−8^; postoperative-day slope in IONX, χ^2^(1) = 9.79, P = 0.00176; group × day interaction, χ^2^(1) = 24.83, P = 6.27 × 10^−7^; Fig. 6g). Retention performance was higher after IONX at PO63, PO70, and PO77 in Holm-adjusted comparisons. A chance-referenced, time-normalized retention area under the curve over the complete PO35–PO70 interval was approximately 1.7-fold greater after IONX (sham, 20.33 ± 3.60, n = 6 mice; IONX, 34.50 ± 1.39, n = 9 mice; two-sided exact permutation test based on the Mann–Whitney U statistic, U = 5.5, P = 0.00879; Fig. 6h). This convergence places durable sensory memory within the same timing-dependent circuit framework that governs learning and the persistence of TC enhancement.

Together, the staggered-training experiments establish reciprocal, state-dependent control between adult TC plasticity and learning. Sensory loss changes when learning is most effective, while learning in turn changes the trajectory of the sensory-loss-induced circuit state. This bidirectional dependence turns timing into a mechanistic variable: the circuit state determines what experience can accomplish, and experience determines how long that state remains available.

## Discussion

Our results identify post-denervation TC plasticity as a reactivated, temporally gated circuit state rather than a static increase in adult cortical responsiveness. This state improves the processing of weak sensory input, lowers perceptual threshold, and promotes learning by accelerating stabilization of learned performance. Its defining property is reciprocal temporal regulation: experience benefits from the state, but also advances or prolongs it according to when training occurs. The failure of identical training to restore potentiation after closure shows that the same experience has different consequences depending on circuit state. Together, these findings shift the interpretation of injury-induced adult plasticity from the presence or absence of plastic potential to its temporal accessibility, establishing state-dependent timing as a principle that links circuit reorganization to adaptive behavior.

The pharmacology links the behavioral advantage of the reactivated state to a molecular mechanism of adult thalamocortical plasticity. Local GluN2B blockade during the early post-denervation period prevented both later L4 response enhancement and accelerated learning, while having little effect on sham responses. This finding extends earlier *ex vivo* evidence that IONX re-engages GluN2B-dependent, silent-synapse-like potentiation at spared TC inputs^21,22^ by demonstrating that reactivation of this critical-period-associated mechanism has a measurable behavioral consequence *in vivo*. More broadly, it shows that sensory loss does not simply alter cortical responsiveness, but can re-engage plasticity mechanisms that increase the capacity of an adult circuit to learn. This provides a mechanistic bridge between reopening of adult plasticity and adaptive behavior. It also recasts critical-period-associated signaling as a functional resource that can be redeployed in the mature brain, providing a general framework for pairing plasticity-promoting mechanisms with appropriately timed training to enhance experience-driven recovery.

The Neuropixels measurements reveal how the reactivated state transforms sensory information at the principal cortical input layer. Enhanced L4 recruitment was concentrated at weak and near-threshold intensities, precisely where behavioral detection improved most strongly, indicating that reactivated TC plasticity preferentially increases the cortical representation of otherwise marginal sensory evidence. At stronger intensities, IONX increased the temporal concentration of evoked spiking, consistent with prior evidence for post-IONX timing synchrony^23^ and established roles for temporal structure in sensory coding.^30,31^ Thus, the reactivated state does not simply amplify cortical activity uniformly; it reshapes sensory coding according to input strength, increasing recruitment when signals are weak while sharpening the temporal structure of responses when signals are strong. This input-dependent transformation provides a circuit-level mechanism through which adult plasticity can expand perceptual access to weak sensory information while preserving precise encoding of stronger inputs. More broadly, these findings show how thalamocortical reorganization can increase the behavioral utility of surviving sensory input after sensory loss. By showing how spared pathways can be retuned to enhance the cortical processing of residual input, the results provide a circuit framework relevant to sensory compensation and to rehabilitation approaches aimed at improving the use of residual sensory signals after peripheral or central injury. These pathway-specific changes complement evidence that behavioral state and movement shape sensory detection and cortical population activity,^32–35^ by identifying how deafferentation retunes the cortical representation of surviving sensory input.

A key conceptual advance is that learning reciprocally reshapes the plasticity state in a phase-dependent manner. Training before the normal peak advanced the emergence of enhanced TC drive, whereas training near or within the active window prolonged it. Unpaired sensory stimulation and reward did not maintain the prolonged state, and training after PO43 did not reinstate potentiation. Thus, experience does not exert a uniform effect on adult plasticity; its impact depends on the circuit state at the time of training. This converts timing from a scheduling variable into a mechanistic component of plasticity. Earlier work showed that spared-whisker sensory experience is required for IONX-induced TC potentiation; trimming the spared whiskers prevented the increase in TC input strength.^22^ Our results extend this observation by showing that the consequences of experience depend on when and under what conditions it engages the plastic circuit: training can advance an emerging state or prolong an active state, whereas training after closure did not reopen the state, and prolonged enhancement depends on sensory–reward contingency. Neuromodulatory and inhibitory systems that gate critical-period plasticity^36–39^ provide candidate pathways through which behavioral salience and task engagement could couple experience to this permissive state. More broadly, these findings provide a circuit-level rationale for why identical training can produce different outcomes when delivered at different stages after neural perturbation, a principle with implications for rehabilitation timing.^40,41^

Learning-associated prolongation was accompanied by more persistent sensory memory, extending the behavioral relevance of the reactivated window beyond acquisition. The convergence of prolonged TC enhancement and persistent behavior places learning and retention within a common temporal framework in which the duration of circuit plasticity and the durability of behavioral change covary. This phase dependence has broader implications for adaptive reorganization after perturbation. Work in peripheral injury has emphasized both maladaptive remapping and large-scale cortical reorganization,^1–11,42^ while studies of experience-dependent plasticity show that training can stabilize pathway- and synapse-specific changes.^43–47^ Our results connect these views by showing that the functional outcome of experience depends on its alignment with an endogenous plasticity trajectory. In this sense, the adult brain is not uniformly plastic after injury; it passes through circuit states with different responsiveness to experience. The logic parallels neurorecovery models in which behavioral intervention interacts with time-limited post-injury plasticity windows,^40,41^ and provides a circuit-level basis for strategies that pair training with periods when surviving pathways are most modifiable.

Together, our findings identify reactivated thalamocortical plasticity as a temporally bounded adult circuit state that both promotes and is modified by learning. The same training can advance an emerging state, prolong an active state, or become ineffective after closure, showing that behavioral outcome depends on alignment with the underlying plasticity trajectory. This reciprocal interaction reveals a circuit-level principle for how adult sensory systems convert transient plastic potential into adaptive change: plasticity establishes a permissive state for learning, and learning in turn shapes the trajectory of that state. More broadly, the findings argue that effective intervention after neural perturbation may depend on matching experience to the state of the circuit rather than applying the same training uniformly across time (Fig. 7).

**Fig. 7.**
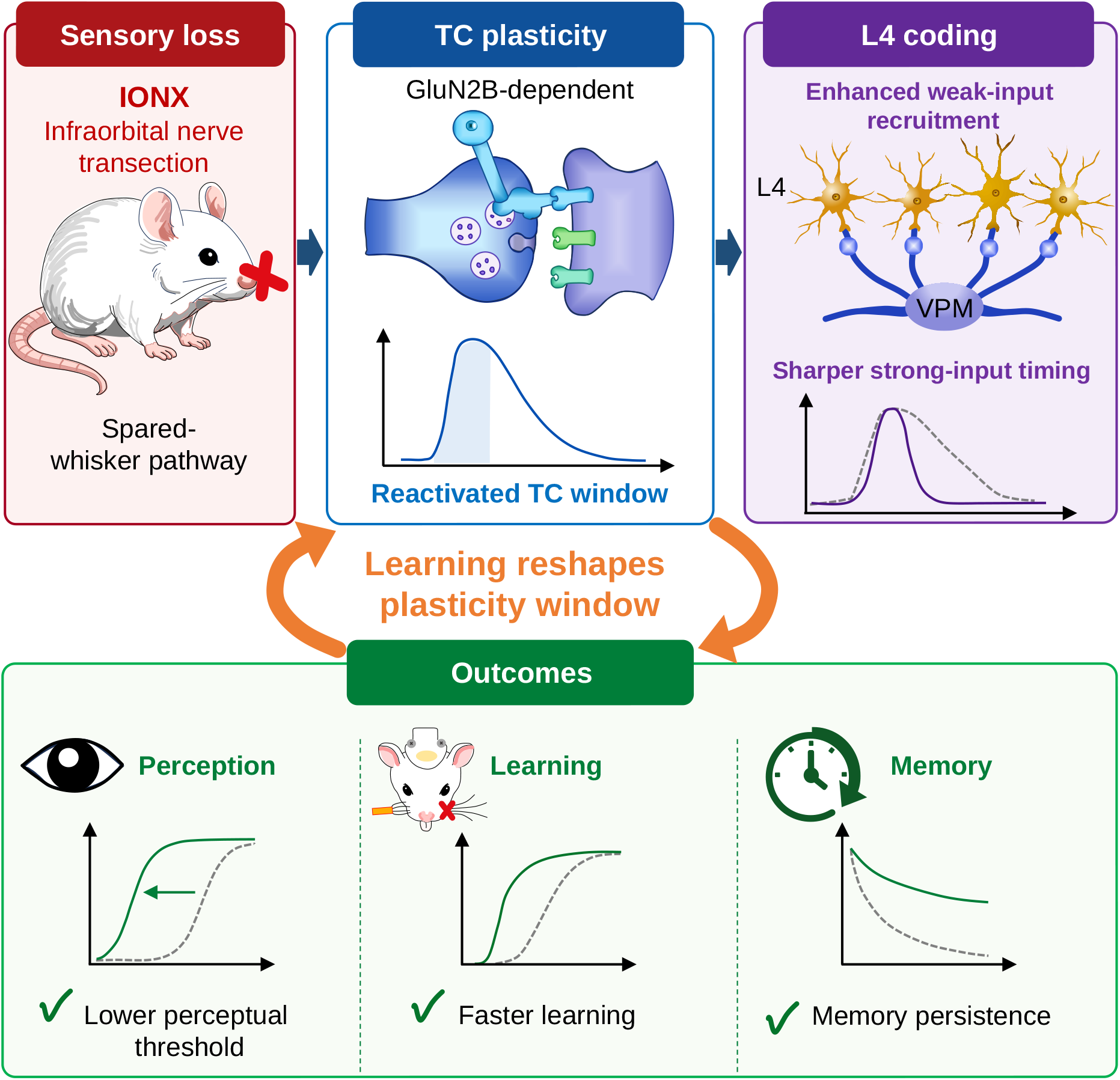
Graphical summary of the reciprocal interaction between reactivated thalamocortical plasticity and experience. Sensory loss reactivates a GluN2B-dependent thalamocortical plasticity window in the spared-whisker pathway. The reactivated state enhances layer 4 coding, lowers perceptual threshold, and promotes faster learning; learning experience then feeds back to reshape the window. Prolongation of the learning-associated window accompanies more persistent sensory memory, linking acquisition and retention within the same timing-dependent framework. Solid green lines indicate IONX and dashed gray lines indicate Sham. Abbreviations: IONX, infraorbital nerve transection.

## Methods

### Animals

Both female and male adult C57BL/6J mice were used. Animals were bred in-house from mice obtained from The Jackson Laboratory and were 8–12 weeks old at surgery. Mice were housed under a 12-h light–dark cycle with food available *ad libitum*; behavioral experiments were performed during the light phase. Mice used for behavior were maintained on a controlled water schedule, with body weight kept at ≥85% of baseline and supplemental water provided as needed. Data were pooled across sex because individual experiments were not designed or powered to test sex-by-surgery or sex-by-treatment interactions. A total of 234 mice were used; experiment-specific analyzed sample sizes are reported in the figure legends and Source Data. All procedures were approved by the Institutional Animal Care and Use Committee under protocol 1160 and followed National Institutes of Health (NIH) and institutional guidelines. Detailed procedures are provided in Supplementary Methods.

### Surgical procedures

For unilateral infraorbital nerve transection (IONX), mice were anesthetized with isoflurane (1–3% in oxygen), the infraorbital nerve bundle was exposed posterior to the whisker pad and transected, and the incision was sutured. Sham mice underwent identical exposure without transection. Ketoprofen (10 mg/kg, intraperitoneal) was administered perioperatively. Postoperative day (PO) denotes time after IONX or Sham surgery. Headplates were implanted under isoflurane anesthesia at least 1 week before behavioral habituation. Contralateral whisker primary somatosensory cortex (wS1) lesions were produced by subpial aspiration of the spared-whisker representation as described previously.^48^ Unless otherwise indicated, analyses were performed in the spared-whisker pathway and corresponding contralateral barrel cortex. Detailed procedures are provided in Supplementary Methods.

### Experimental design and whisker-detection behavior

Separate cohorts were used for postoperative physiology, pharmacology, behavioral acquisition, Neuropixels recordings, learning-timing experiments, unpaired controls, and memory-retention assays. Head-fixed mice performed a Go/No-go whisker-detection task controlled in LabVIEW. A calibrated coil-driven actuator delivered 2-ms deflections to whiskers inserted approximately 4 mm from the pad; 70-dB white noise masked actuator-associated sounds.^23^ Sessions comprised 100 trials. Go/No-go trial ratios were 90:10 in session 1, 70:30 in session 2, and 50:50 thereafter. Trial initiation required a randomized 2.5–3.5-s no-lick period. Licking within 1 s after a Go stimulus was scored as a hit and rewarded with approximately 10 µL water; licking during the corresponding No-go window was scored as a false alarm. Intertrial intervals were 3–4 s initially and were extended to 6–12 s when impulsive licking increased.

Primary acquisition began at PO17 with a 3.3° deflection. Criterion was ≥80% hit rate and ≤30% false-alarm rate for two consecutive sessions, with training limited to 21 sessions. Independent cohorts were trained at fixed amplitudes of 1.8°, 3.3°, or 8.3°. Sequential-training cohorts progressed through 3.3°, 2.5°, and 1.8° before retention or Neuropixels threshold testing, following an established psychometric design.^24^ At 2.5° and 1.8°, the same performance thresholds were required for one session, with a six-session limit at each intensity. Learning-timing cohorts began 3.3° training at PO4, PO10, PO17, or PO43. Post-learning and pre-training wS1 lesions and trained-whisker removal were used to test dependence on barrel cortex and whisker-mediated input. Detailed apparatus, calibration, training, and control procedures are provided in Supplementary Methods.

### Learning-state and motor-output analyses

Trial-by-trial behavior was segmented with a three-state left-to-right hidden Markov model (HMM) representing pre-learning, active-learning, and post-learning states.^26–29^ Daily sessions were divided into non-overlapping 20-trial blocks, and hit and false-alarm counts were modeled as state-specific binomial emissions. Hit probability was constrained to increase across states, false-alarm probability to be higher in pre-learning than in post-learning, and backward transitions were disallowed. Parameters were estimated by Baum–Welch expectation–maximization and state sequences decoded with the Viterbi algorithm. Learning onset and completion were defined as the first trials in the first active-learning and post-learning blocks, respectively. Completers reached post-learning and had pooled post-learning hit rate ≥0.65 and false-alarm rate ≤0.35; a predefined plateau rule captured fast learners who reached ceiling before a decoded post-learning transition. Full model specification, initialization, convergence criteria, and plateau rule are provided in Supplementary Methods.

Group differences in completer fraction were assessed with Fisher’s exact test; onset, completion, and the onset-to-completion interval were compared with two-sided Mann–Whitney U-tests. State-specific fitted hit and false-alarm probabilities were logit-transformed and analyzed with linear mixed-effects models containing group, state, and their interaction, with a random intercept for mouse. Lick latency, inter-lick interval, and bout duration were summarized by mouse and HMM state and analyzed with analogous mixed-effects models; latency and inter-lick interval were log-transformed. Holm correction was applied within prespecified contrast families. Detailed model-selection procedures are provided in Supplementary Methods.

### Memory retention and unpaired control

After sequential training, retention cohorts were probed weekly at 3.3° for approximately 2 months using 20-trial sessions (10 Go and 10 No-go trials) to minimize relearning. Retention was analyzed with a Gaussian generalized estimating equation model containing group, centered postoperative day, and their interaction, with mouse as the clustering variable. A complementary chance-referenced, time-normalized area under the curve was calculated for PO35–PO70 and compared by an exact permutation test based on the Mann–Whitney U statistic.

To test whether prolonged TC enhancement required sensory–reward contingency, unpaired-control mice received six daily 100-trial sessions containing 50 stimulus and 50 no-stimulus trials. Forty water rewards were delivered independently at random times during intertrial intervals and never within the response window. All other task conditions matched paired training. Whisker-evoked L4 responses were measured at PO70. Full retention and unpaired-control procedures are provided in Supplementary Methods.

### Ro25-6981 pharmacology

Ro25-6981 maleate (1 µM in sterile phosphate-buffered saline (PBS); Tocris Bioscience) or vehicle was delivered locally to left wS1 with osmotic minipumps (model 2002; Alzet) connected to an L4-targeted cannula (anteroposterior (AP), −1.5 mm; mediolateral (ML), 3.2 mm; dorsoventral (DV), −0.5 mm from the cortical surface). Infusion began at IONX or Sham surgery and proceeded at approximately 0.5 µL/h for 14 days; pumps were removed on PO21. This design targeted the early induction phase of GluN2B-dependent TC plasticity rather than acute drug effects during behavior.^22^ Training began at PO17. Four groups were studied (Sham + Vehicle, n = 9; Sham + Ro25-6981, n = 8; IONX + Vehicle, n = 9; IONX + Ro25-6981, n = 9). Cannula placement was verified histologically, and Alexa Fluor 488–dextran infusion using the same configuration estimated delivery spread. Animals were excluded when the cannula track did not overlap wS1 barrel cortex. In learners, post-training TC responses were recorded at age-matched postoperative times between PO33 and PO70. Additional surgical and validation details are provided in Supplementary Methods.

### Electrophysiology and spike analysis

Conventional 32-channel silicon-probe recordings were used in urethane-anesthetized mice to measure VPM and L4 local field potentials and population spiking across postoperative time and after pharmacology. A fixed 6.7° whisker deflection was used for these assays. Simultaneous VPM–L4 recordings targeted barrel cortex (AP, −1.5 mm; ML, 3.2 mm; DV, −1.25 mm; 30° insertion) and VPM (AP, −1.8 mm; ML, 1.45 mm; DV, −4.0 mm). Signals were acquired at 30 kHz.

Neuropixels 1.0 probes^49^ recorded wS1 and VPM activity during behavioral threshold testing after criterion, between PO33 and PO70. Catch trials and 14 nonzero deflections (0.3°, 0.4°, 0.5°, 0.6°, 0.7°, 0.8°, 0.9°, 1.0°, 1.3°, 1.8°, 3.3°, 5.5°, 8.3°, and 11°) were pseudorandomly interleaved, with 20 trials per condition. Data were processed with Kilosort4^50^ and curated in Phy. Well-isolated “good” units formed the single-unit dataset; multi-unit activity was additionally included for population recruitment analyses. L4 and VPM sensory responses were quantified over 0–50 ms and 0–20 ms after stimulus onset, respectively, relative to duration-matched prestimulus baselines. For group inference, validated activity sites were averaged within each mouse at each intensity, and mouse was the biological replicate. Acquisition, targeting, and quality-control details are provided in Supplementary Methods.

### LFP and current-source-density analyses

LFP signals were downsampled to 1 kHz, band-pass filtered at 0.1–300 Hz, baseline-corrected over −10 to 0 ms, and averaged across trials. Evoked peak amplitude was the magnitude of the largest negative-going deflection within 0–50 ms in L4 or 0–20 ms in VPM. Laminar assignments were anchored to probe depth and the principal early whisker-evoked current sink. Current-source density (CSD) was calculated from the second spatial derivative of the depth-aligned LFP after smoothing across depth and time; positive values were interpreted as sinks. Histology and depth mapping supported L4 assignment.

### Perceptual-threshold and temporal-sharpening analyses

For each mouse, lick-response probability was related to mouse-level mean stimulus-evoked L4 Δ spikes per trial across intensities. Neural and behavioral coordinates were independently range-scaled, and each admissible internal transition was evaluated with a two-segment total least-squares model. The minimum-error transition was accepted as the operational perceptual threshold when the post-transition behavioral slope exceeded the pre-transition slope. Fitting was restricted to intensities ≤1.8° to isolate the low-intensity transition and prevent near-asymptotic responses from dominating the fit. Group differences were assessed with a two-sided exact permutation test based on the Mann–Whitney U statistic. Manual transition estimates provided a sensitivity analysis.

The temporal sharpening index (TSI) was the fraction of sensory-evoked spikes contained in the peak 10-ms window within 0–100 ms after stimulus onset, consistent with established roles of temporal response structure in sensory coding.^30,31^ The peak window was selected separately for each unit and intensity. Unit-level TSI values were averaged within mouse. ΔTSI was mean TSI at high intensities (3.3°, 5.5°, 8.3°, and 11°) minus mean TSI at low intensities (0.3°, 0.4°, and 0.5°), and groups were compared at the mouse level.

### Histological verification

Recording sites, probe trajectories, drug-delivery sites, and laminar landmarks were verified after perfusion. VPM barreloids and wS1 barrels were visualized by cytochrome oxidase histochemistry and fluorescence imaging. For VPM barreloid mapping, brains were sectioned in an oblique plane guided by prior anatomical descriptions;^51^ wS1 cortex was sectioned tangentially through L4. An adeno-associated virus serotype 9 (AAV9) vector expressing CaMKIIα-hChR2(E123A)-mCherry^52^ was used in validation animals to delineate VPM axons in the wS1 barrel field. DiI-labeled probe tracks, anatomical maps, and CSD landmarks were combined to confirm inclusion. Detailed histological procedures are provided in Supplementary Methods.

### Randomization, blinding, and exclusion criteria

Group assignment balanced litter, sex, and surgery date. Investigators were necessarily aware of Sham and IONX procedures, but analyses used predefined criteria and coded identifiers when feasible. Exclusions were limited to technical failures, unsuccessful recovery, unconfirmed targeting, or incomplete data caused by technical interruption.

### Statistics and reproducibility

Statistical analyses were performed in Prism, R, and Python. Two-group comparisons used parametric or nonparametric tests according to design and distribution; multi-group comparisons used ANOVA or nonparametric alternatives with appropriate multiplicity correction. Repeated trajectories were analyzed with repeated-measures, mixed-effects, or generalized estimating equation models. Time-to-criterion analyses used log-rank Mantel–Cox tests with non-learners censored. Psychometric and stimulus-response curves were analyzed with mixed ANOVA or mixed-effects models, followed by Holm-adjusted comparisons across intensities. Mouse identity defined the biological replicate for repeated and Neuropixels analyses. Unless otherwise stated, data are mean ± SEM. Exact P values, sample sizes, and effect estimates are reported in figure legends and Source Data. No statistical method was used to predetermine sample size; sample sizes were based on prior studies with similar designs. Detailed outcome-specific analyses are provided in Supplementary Methods.

## Data availability

Source data are provided with this paper. Additional processed behavioral and electrophysiological datasets, together with per-animal metadata supporting the figures, will be made available to editors and reviewers during peer review and deposited in a publicly accessible, DOI-minting repository before publication. Repository accession information will be added to the final manuscript.

## Code availability

Custom Python and R code used for HMM-based behavioral modeling, Neuropixels analyses, and statistical testing is provided as a reviewer-accessible submission file and will be deposited in a publicly accessible, DOI-minting repository before publication. Repository accession information and software-environment specifications will be added to the final manuscript.

## Acknowledgements

We thank Soohyun Lee and Emily Petrus for critical reading of the manuscript and helpful comments. We thank Nadia Bouraoud for technical assistance and George Dold and the Section on Instrumentation for engineering support.

## Funding

This research was supported by the Intramural Research Program of the National Institutes of Health (NIH). The contributions of the NIH author(s) were made as part of their official duties as NIH federal employees, are in compliance with agency policy requirements, and are considered Works of the United States Government. However, the findings and conclusions presented in this paper are those of the author(s) and do not necessarily reflect the views of the NIH or the U.S. Department of Health and Human Services.

## Author contributions

Conceptualization, H.J. and A.P.K.; methodology, H.J. and A.P.K.; software, H.J. and Z.-D.D.; formal analysis, H.J. and Z.-D.D.; investigation, H.J. and N.P.; resources, K.S. and A.P.K.; data curation, H.J.; writing – original draft, H.J.; writing – review & editing, H.J., N.P., Z.-D.D., K.S., and A.P.K.; visualization, H.J. and N.P.; supervision, A.P.K.; project administration, H.J. and A.P.K.; funding acquisition, A.P.K.

## Competing interests

The authors declare no competing interests.

## Notes

### Competing Interest Statement

The authors have declared no competing interest.

