## Supplementary Information for "A reactivated thalamocortical plasticity window promotes learning and is reshaped by experience"

Contents: Supplementary Methods; Supplementary Figs. 1–8; Supplementary References; Supplementary Table 1.

### Supplementary Methods

The following sections provide detailed procedures and analytical specifications that complement the concise Methods in the main manuscript.

#### Unilateral sensory deafferentation

Mice underwent unilateral infraorbital nerve transection (IONX) under isoflurane anesthesia (1–3% in oxygen). Ophthalmic ointment was applied, and body temperature was maintained at approximately 37 °C. A 1–2-mm incision was made immediately posterior to the whisker pad to expose the infraorbital nerve bundle. In IONX mice, the nerve bundle was transected with fine scissors and, where required, the cut ends were cauterized using a fine low-temperature cautery tip (H100; Bovie, Aspen Surgical). Sham-operated mice underwent the same anesthesia, incision, and nerve exposure, but the nerve was left intact. The incision was closed with sutures, ketoprofen (10 mg/kg, intraperitoneal) was administered perioperatively, and animals were recovered on a warming surface and monitored daily.

#### Headplate implantation

For head-fixed behavioral experiments, mice were implanted with a custom headplate designed in SOLIDWORKS 3D CAD software (Dassault Systèmes) and fabricated by stereolithography using a ProJet 6000 HD printer and Accura ClearVue photopolymer resin (3D Systems). Mice were anesthetized with isoflurane (1–3% in oxygen), positioned in a stereotaxic frame, and maintained at approximately 37 °C using a heating pad. Ophthalmic ointment was applied to prevent corneal drying. After the scalp was removed, the skull surface was cleaned and dried, and the headplate was positioned and secured using C&B Metabond Quick Adhesive

Cement (Parkell), prepared according to the manufacturer's instructions. Ketoprofen (10 mg/kg, intraperitoneal) was administered perioperatively for analgesia. Mice were returned to their home cages after recovery from anesthesia and allowed to recover for at least 1 week before water scheduling and head-fixation habituation.

**Cortical lesion of contralateral whisker primary somatosensory cortex**

Contralateral whisker primary somatosensory cortex (wS1) lesions were performed under aseptic surgical conditions. Mice were anesthetized with isoflurane (1–3% in oxygen), placed in a stereotaxic frame, and maintained on a heating pad throughout surgery. The wS1 region corresponding to the spared-whisker representation was identified using a mouse brain atlas. For lesions in the left hemisphere, the craniotomy boundary was delineated by connecting multiple atlas-based stereotaxic landmarks spanning wS1: anteroposterior (AP), +0.5 to −2.1 mm, and mediolateral (ML), 2.2–4.5 mm relative to bregma. A craniotomy was made along this outline, and the dura was carefully opened to expose the cortical surface.

Cortical tissue within the spared-whisker representation was removed by manual subpial aspiration, as described previously.[1] Briefly, a sterile blunt-tipped needle connected to a syringe was used to apply gentle, hand-controlled negative pressure. Cortical gray matter was aspirated gradually in small steps under visual guidance, with the aspiration depth limited to the cortical mantle. Aspiration was stopped when the underlying white matter became visible, and care was taken to avoid damage to subcortical structures. The lesion cavity and cortical surface were rinsed with sterile phosphate-buffered saline (PBS), the scalp was closed, and mice were allowed to recover on a heating pad before returning to their home cage.

**Experimental design**

Separate cohorts were used for postoperative time-course electrophysiology, pharmacology, behavioral acquisition, Neuropixels recordings, timing-dependent training, unpaired controls, and memory-retention assays, as specified in the figure legends and Source Data. Whisker-evoked responses were measured in layer 4 (L4) barrel cortex across postoperative days to define the time course of thalamocortical (TC) enhancement. Behavioral training was initiated at defined postoperative time points to test how TC state influenced learning and how learning altered the TC plasticity window. Local GluN2B blockade, Neuropixels recordings, unpaired controls, and retention assays were used to test molecular necessity, circuit-level correlates, experience dependence, and memory persistence, respectively. Key reagents, software, and instrumentation are listed in Supplementary Table 1.

**Behavioral apparatus and whisker stimulation**

Behavioral training was conducted in head-fixed mice performing a Go/No-go whisker-detection task. Task control, stimulus timing, lick detection, reward delivery, and trial-by-trial behavioral data acquisition were implemented in LabVIEW 2019 SP1 (National Instruments) and interfaced through an NI USB-6212 multifunction I/O device. To minimize electrical noise during *in vivo* electrophysiological recordings, the headplate-coupling component of the restraint assembly was machined from electrically insulating G-10 glass-fiber-reinforced epoxy laminate (Garolite) and secured with polyether ether ketone (PEEK) screws. Whisker stimuli were delivered using a custom coil-driven actuator adapted from our previously characterized whisker-stimulation system[2] and validated for the present experiments (Supplementary Fig. 2a,b). The actuator was driven by a pulse generator and audio amplifier and consisted of a capillary glass tube attached to a custom voice coil. During stimulation, the whiskers were inserted into the capillary tube approximately 4 mm from the whisker pad. Each stimulus consisted of a 2-ms whisker deflection at a defined angular amplitude.

Actuator displacement and timing were calibrated using a laser displacement sensor (LK-H022, Keyence; repeatability, 0.02 µm; Supplementary Fig. 2c). Command voltage was converted to displacement using the laser-calibrated voltage–displacement relationship, and angular deflection was calculated from the measured displacement and the distance between the whisker pad and the stimulation point. Background white noise (70 dB) was presented throughout behavioral sessions to minimize actuator-associated auditory cues. Before formal training, mice were habituated to head fixation, water scheduling, lick training, and whisker insertion.

**Whisker stimulus amplitudes and acquisition training**

Behavioral sessions consisted of 100 trials. To facilitate acquisition, Go and No-go trials were presented in pseudorandom order at ratios of 90:10 in the first session, 70:30 in the second session, and 50:50 from the third session onward. The intertrial interval was 3–4 s during early training and was extended to 6–12 s when impulsive licking increased. Following the intertrial interval, trial initiation required a randomized 2.5–3.5-s no-lick baseline; licking during this period aborted the trial. On Go trials, a whisker deflection was delivered at trial onset, and licking within the subsequent 1-s response window was scored as a hit and triggered an approximately 10-µL water reward; failure to lick was scored as a miss. On No-go trials, no stimulus was delivered, and licking or withholding licking during the response window was scored as a false alarm or correct rejection, respectively. During training, mice received *ad* *libitum* water for 1–2 days per week and were monitored regularly for body weight, hydration, and general health under veterinary oversight.

For the primary acquisition experiments, mice began training on postoperative day 17 (PO17) with a 3.3° whisker deflection. This amplitude was selected to preserve sensitivity to group differences: it reliably supported task acquisition in Sham mice, was lower than the 6.7° deflection used for postoperative time-course physiological measurements, and lay within the dynamic, non-saturating range of the actuator input–output relationship. Learning criterion was defined as a hit rate of ≥80% and a false-alarm rate of ≤30% for two consecutive sessions, with sessions to criterion serving as the primary acquisition measure unless otherwise indicated. To determine how stimulus strength influenced learning, independent cohorts were trained at fixed deflection amplitudes of 1.8°, 3.3°, or 8.3°.

For the PO17 sequential-training cohorts used for retention or perceptual-threshold testing, mice were trained successively at 3.3°, 2.5°, and 1.8°. This progression was adapted from an established psychometric whisker-detection paradigm in which animals advanced from a readily detectable stimulus to weaker stimuli before randomized multi-intensity testing.[3] Criterion at 3.3° required a hit rate of ≥80% and a false-alarm rate of ≤30% for two consecutive sessions, with training limited to 21 sessions. At 2.5° and 1.8°, mice advanced after meeting the same thresholds in a single session, with training limited to six sessions at each intensity. After reaching criterion, one subset underwent weekly retention testing at 3.3°, whereas another completed a single Neuropixels threshold-testing session, as described below.

**Post-learning and pre-training wS1 lesion experiments**

Two lesion paradigms were used to assess the requirement of contralateral wS1 for whisker-detection behavior. For post-learning lesions, Sham and IONX mice were trained from PO17 with a 3.3° whisker deflection until they reached learning criterion. Contralateral wS1 representing the spared-whisker pathway was lesioned on the day after criterion was reached. After a one-day recovery period, mice were returned to the same Go/No-go task and retrained for four consecutive days. For pre-training lesions, contralateral wS1 was lesioned before PO17 training onset. After recovery, Sham and IONX mice were trained from PO17 using the same Go/No-go protocol and learning criterion as intact mice. Mice that did not reach criterion within the training period were classified as non-learners.

**Whisker-trimming control experiment**

To determine whether task performance depended on whisker-mediated tactile input, a subset of trained mice was tested after removal of the trained whiskers. After reaching criterion, the trained whiskers were trimmed to the base under brief isoflurane anesthesia (1–2% in oxygen) in both the 3.3° and 8.3° cohorts. Following recovery, mice were tested in an otherwise identical behavioral session using the same trial structure, intertrial intervals, no-lick baseline periods, actuator command voltages, and background white-noise masking as during training. Including the 8.3° cohort allowed us to test whether performance at a suprathreshold stimulus intensity also required direct whisker contact, thereby excluding the possibility that animals could rely on actuator-associated auditory, vibrational, or other non-tactile cues even under strong-stimulus conditions. This design preserved the non-tactile sensory context while selectively eliminating whisker-mediated tactile input.

### Behavioral control analyses

First-day lick probability was calculated as the percentage of all trials containing a lick during the 1-s response window in the first acquisition session. High lickers were defined as mice with a first-day lick probability greater than 50%. These analyses pooled paired-training cohorts and excluded unpaired-control and wS1-lesion cohorts. The association between first-day lick probability and sessions to criterion was evaluated using descriptive linear regression; mice that did not reach criterion within 21 sessions were assigned a plotting value of 22 sessions for this analysis. High-licker fractions were compared using a two-sided Fisher's exact test, and sessions to criterion were compared using two-sided Mann–Whitney U-tests.

### Hidden Markov modeling of learning-state transitions

Trial-by-trial behavioral performance was analyzed using a three-state left-to-right hidden Markov model (HMM) to segment each mouse's training trajectory into pre-learning, active-learning, and post-learning states. The three-state structure was chosen *a priori* to separate pre-learning performance from the active transition period and the stable post-learning state; this avoided collapsing the steep rise in discrimination into either baseline or asymptotic performance and allowed learning onset and learning completion to be quantified separately. Each 100-trial daily session was divided into non-overlapping 20-trial blocks. Terminal fragments containing fewer than 10 valid trials were merged into the preceding block, whereas larger terminal fragments were retained. For each block, the observed data consisted of the number of hits among Go trials, false alarms among No-go trials, and the corresponding number of Go and No-go trials. For each block, emissions were modeled as two independent binomial observations conditional on latent state: one for hits on Go trials and one for false alarms on No-go trials. For a block with hit and false-alarm counts ($n_{\text{hit}}$, $n_{\text{FA}}$) out of $n_{\text{Go}}$ Go trials and $n_{\text{NoGo}}$ No-go trials, the emission log-likelihood in latent state $k$ was

$$\log p(y_b \mid z_b = k) = \log \text{Bin}\left(n_{\text{hit},b};\ n_{\text{Go},b}, p_{\text{hit},k}\right) + \log \text{Bin}\left(n_{\text{FA},b};\ n_{\text{NoGo},b}, p_{\text{FA},k}\right), \tag{1}$$

where $y_b$ is the observed data for block $b$; $z_b$ is the latent state of block $b$, which takes one of three values $k \in \{0,1,2\}$ corresponding to the pre-learning, active-learning, and post-learning states. $p_{\text{hit},k}$ and $p_{\text{FA},k}$ denote the state-specific hit and false-alarm probabilities, respectively.

State-specific emission probabilities were constrained so that hit probability increased monotonically across states, whereas false-alarm probability was higher before learning than after learning. These constraints were enforced at each maximization step: after the closed-form binomial update, hit probabilities were projected onto the monotonically increasing ordering using the pool-adjacent-violators algorithm for weighted isotonic regression, and a minimum separation of 0.05 was imposed between adjacent states so that the three states remained distinct. Emission probabilities were initialized from the 25th, 50th, and 75th percentiles of the smoothed block-wise hit and false-alarm rates, with the false-alarm percentiles assigned in descending order so that the pre-learning state began with the highest false-alarm rate. The transition model was left-to-right, with backward transitions forbidden. Separate transition matrices were used for consecutive blocks within the same session and for transitions across sessions. The initial state distribution was uniform across the three states.

Model parameters were estimated with the Baum–Welch expectation–maximization algorithm.[4,5] The expectation step computed block-level posterior state probabilities and two-state transition marginals using the forward–backward algorithm, applying the within-session or between-session transition matrix at each block boundary. The maximization step updated binomial emission probabilities and transition probabilities in closed form. Expectation–

maximization was run for up to 300 iterations and terminated when the change in marginal log-likelihood fell below $10^{-5}$ or when a predefined rolling limit-cycle criterion was reached. Because the maximization step included order and minimum-separation constraints on state-specific emission probabilities, convergence was assessed using the termination criteria described above rather than by assuming unconstrained Baum–Welch monotonicity.

After parameter estimation, the most probable block-level state sequence was decoded using the Viterbi algorithm.[4] The decoded sequence was constrained to be non-decreasing in state index, so that once a mouse entered a later learning state it could not return to an earlier state (Fig. 2a). This encoded the assumption that learning-state progression is irreversible (pre-learning → active-learning → post-learning). Each trial was assigned the state label of its 20-trial block. Learning onset was defined as the first trial in the first active-learning block, and learning completion as the first trial in the first post-learning block. Mice were classified as HMM-defined learning completers if they reached the post-learning state and pooled post-learning performance met a quality gate of hit rate ≥0.65 and false-alarm rate ≤0.35. This gate differed from the behavioral acquisition criterion because it evaluated pooled performance within an inferred state rather than criterion attainment across consecutive sessions; the symmetric 0.65/0.35 cutoffs accommodated block-level variability while requiring stimulus-selective responding rather than generalized licking. Mice that entered the active-learning state but did not reach post-learning, or reached post-learning but failed the performance criteria, were classified as learning incomplete. Mice that never left pre-learning were classified as non-learners. For completers, the consolidation interval was the number of trials between learning onset and completion. To avoid misclassifying fast learners whose performance reached ceiling before the session ended (and before the Viterbi decoder could reach the post-learning transition), a mouse initially classified as learning incomplete was reclassified as a completer if its active-learning emission estimates satisfied the same hit-rate and false-alarm-rate criteria and its decoded active-learning blocks contained a run of at least three consecutive blocks with an observed hit rate at or above the hit-rate threshold. For these reclassified mice, learning completion was assigned to the first block of the qualifying three-block plateau.

Group differences in the fraction of HMM-defined learning completers were assessed using Fisher's exact test. Learning onset, learning completion, and consolidation interval were compared between groups using two-sided Mann–Whitney U-tests because trial-count distributions were right-skewed. For HMM-defined completers, state-specific task performance was quantified using each mouse's fitted HMM emission probabilities for hits on Go trials and false alarms on No-go trials. Because these probabilities were bounded between 0 and 1, each was logit-transformed before analysis. Each transformed measure was analyzed using a linear mixed-effects model in R (nlme package), with fixed effects of group (Sham, IONX), learning state, and their interaction, and a random intercept for mouse. Fixed effects were assessed using Type III F-tests. For each measure, a homoscedastic model was compared with a model allowing state-specific residual variances (varIdent), and the final model was selected by Bayesian information criterion. Estimated marginal means and pairwise contrasts were computed using the emmeans package and back-transformed to proportions. Learning-state and group contrasts are reported as odds ratios, and P values for families of state-wise contrasts were Holm-adjusted.

**Licking motor-output analysis**

Licking behavior was characterized by three measures: lick latency, inter-lick interval (ILI), and bout duration. Lick latency was defined as the time from stimulus onset to the first lick within the 1-s post-stimulus response window. ILI was calculated as the mean interval between consecutive licks within a bout and was defined for hit trials with at least two licks. Bout duration was defined as the time from the first to the last lick within a bout. Analyses were restricted to HMM-defined learning completers (Sham, n = 19 mice; IONX, n = 26 mice). All available state-specific observations were retained, and the mixed-effects models accommodated missing state-specific measurements. For each mouse, the median of each measure was computed within each HMM-defined learning state.

Each licking variable was analyzed using a linear mixed-effects model in R (nlme package), with fixed effects of group (Sham, IONX), learning state (pre-learning, active learning, post-learning), and their interaction, and a random intercept for mouse to account for repeated measurements. Fixed effects were assessed with Type III F-tests. Before model fitting, each measure was examined for skewness and for state-dependent changes in within-subject variance. Lick latency and ILI were strictly positive and right-skewed and were therefore log-transformed to improve residual normality. Bout duration was approximately symmetric on the raw scale and was analyzed without transformation. For each measure, a homoscedastic model was compared with a model allowing state-specific residual variances (varIdent), with model selection based on Bayesian information criterion. The state-specific variance model was strongly preferred for lick latency, whereas homoscedastic models were retained for ILI and bout duration. Residual and quantile–quantile plots confirmed approximately normal residuals under the selected models. Estimated marginal means and pairwise contrasts were computed using the emmeans package. Pre-planned contrasts included consecutive learning-state comparisons within each group, group comparisons within each learning state, and group × state interaction contrasts. P values were adjusted using the Holm method.

**Weekly memory-retention assay**

After mice in the PO17 retention subset completed sequential criterion-based training at 3.3°, 2.5°, and 1.8°, memory retention was assessed at 3.3° once per week for approximately 2 months using sparse probe sessions designed to minimize relearning. The sequential training procedure was adapted from a previously established psychometric whisker-detection paradigm[3] and was used to establish stable detection performance across progressively weaker stimulus intensities before retention testing.

Each weekly probe session consisted of 20 trials, including 10 Go trials and 10 No-go trials presented in pseudorandom order. Weekly probe performance was used to quantify memory persistence and to derive the retention summary metrics specified in the figure legends. Retention performance was analyzed using a Gaussian generalized estimating equation (GEE) with an exchangeable working correlation and robust covariance. Percent correct was modeled as a function of group, centered postoperative day, and their interaction, with mouse identity specified as the clustering variable. The model included all available observations at the eight scheduled postoperative time points common to both groups from PO35 to PO84. Postoperative day was centered at PO59.5. Sham mice were assessed throughout PO35–PO84 ($n = 6$), whereas IONX mice were assessed through PO70 ($n = 9$), with four mice remaining at PO77 and PO84. The reduction in IONX sample size after PO70 resulted from planned terminal thalamocortical physiology experiments and was unrelated to behavioral performance; no missing values were imputed. Between-group comparisons at each scheduled retention time point were performed using two-sided Welch's t-tests, with P values adjusted across the eight comparisons using the Holm method.

As a complementary individual-level summary, a chance-referenced, time-normalized retention area under the curve (AUC) was calculated for each mouse over PO35–PO70, the complete observation interval available for all included animals. Percent-correct performance was referenced to the 50% chance level and integrated across postoperative days using the trapezoidal rule. Values below chance were retained as negative contributions. The resulting AUC was divided by the 35-day observation interval, yielding the mean percentage-point deviation from chance across PO35–PO70. Group differences were evaluated using a two-sided exact permutation test based on the Mann–Whitney U statistic.

**Unpaired control**

To determine whether maintenance of the potentiated TC state required associative learning rather than repeated exposure to sensory stimulation and reward, mice underwent one unpaired-control session per day for six consecutive days. Each session consisted of 100 trials, comprising 50 whisker-stimulus trials and 50 no-stimulus trials presented in pseudorandom order using the same trial structure as in the paired Go/No-go task. On stimulus trials, the whiskers were deflected by 3.3°.

Forty water rewards were delivered per session, approximating the number typically obtained after criterion-level performance in the paired task. Reward delivery was independent of trial type, whisker stimulation, and licking. Rewards were delivered at randomly selected times during the 7–12-s intertrial intervals and never occurred within the 1-s response window after trial onset. Thus, licking did not trigger reward delivery, and rewards did not reliably follow either licking or whisker stimulation, preventing formation of a stimulus–response–reward association.

All other experimental conditions, including head fixation, trial number, the proportions of stimulus and no-stimulus trials, whisker-stimulus intensity, intertrial intervals, response-window duration, behavioral apparatus, and background white-noise masking, were matched to those used in the paired-training condition. Whisker-evoked L4 responses were subsequently recorded at PO70 to determine whether non-associative exposure to whisker stimulation and reward alone was sufficient to maintain the IONX-induced potentiated TC state.

**Ro25-6981 pharmacology**

Ro25-6981 maleate (Tocris Bioscience), a selective antagonist of GluN2B-containing N-methyl-D-aspartate (NMDA) receptors, was delivered locally to wS1 using osmotic minipumps (model 2002; Alzet). Ro25-6981 was freshly prepared in sterile PBS at a final concentration of 1 μM, and sterile PBS was used as the vehicle. Pumps were filled with 200 μL of solution, primed overnight in sterile PBS at 37 °C, and connected to an infusion cannula targeted to L4 of the left wS1 (AP, −1.5 mm; ML, 3.2 mm; dorsoventral (DV), −0.5 mm from the cortical surface).

IONX or Sham surgery, pump and cannula implantation, and headplate implantation were performed during the same surgical session under isoflurane anesthesia (1–3% in oxygen), such that infusion began immediately after surgery. Model 2002 pumps delivered solution at approximately 0.5 μL/h for 14 days, corresponding to an estimated Ro25-6981 delivery rate of 0.5 pmol/h. Pumps were removed from all animals on PO21, after the first four days of behavioral training, to avoid surgery immediately before training onset.

The treatment was designed to inhibit GluN2B-dependent thalamocortical plasticity during the early postoperative induction window rather than to test acute drug effects during behavior. This interval encompassed the previously reported increase in GluN2B after IONX, beginning around PO9, peaking at PO12, and returning to baseline by PO15.[6]

In a separate validation experiment, Alexa Fluor 488-conjugated dextran (molecular weight 10,000; anionic and fixable; Invitrogen, Thermo Fisher Scientific) was infused using the same pump and cannula configuration to estimate the spatial extent of delivery. Brains were fixed, coronally sectioned, and imaged for dextran fluorescence. Because the dextran diffusion field did not encompass the entire barrel cortex, peripheral whiskers were trimmed before behavioral testing and stimulation was restricted primarily to centrally located principal whiskers whose cortical representations overlapped the observed dextran infusion field. In Ro25-6981- and vehicle-infused animals, cannula tracks were examined histologically, and animals were excluded if the track did not overlap the intended wS1 barrel cortex target.

Four behavioral pharmacology groups were studied (n = 35 mice total): Sham + Vehicle (n = 9), Sham + Ro25-6981 (n = 8), IONX + Vehicle (n = 9), and IONX + Ro25-6981 (n = 9). Cannula-tip locations were histologically verified across these animals (Supplementary Fig. 6c). Behavioral experiments tested whether early GluN2B blockade altered the subsequent IONX-associated acceleration of whisker-detection acquisition. In mice that reached criterion, whisker-evoked TC responses were recorded between PO33 and PO70, a postoperative interval selected to assess the sustained phase of learning-associated TC potentiation in IONX mice. Sham and IONX mice within each cohort were recorded within 1–2 days of one another. Because IONX mice often reached criterion earlier, recording was delayed until the corresponding Sham mice completed training to permit postoperative age-matched comparisons.

**_In vivo_ electrophysiology**

*In vivo* electrophysiology was used to measure whisker-evoked local field potentials and population spiking. Conventional 32-channel NeuroNexus silicon-probe recordings were performed in urethane-anesthetized mice to define the postoperative time course of TC potentiation after IONX, whereas high-density Neuropixels 1.0 recordings were performed during behavioral threshold testing to relate L4 spiking to perceptual performance. The postoperative ages used for conventional silicon-probe recordings are specified in Supplementary Fig. 1.

For conventional silicon-probe recordings, whisker-evoked responses were recorded from the ventral posteromedial thalamic nucleus (VPM) and barrel cortex L4 using 32-channel probes (A1x32-6mm-50-177-A32; NeuroNexus Technologies). For postoperative time-course and pharmacological recordings, TC responses were evoked using a fixed 6.7° whisker deflection, corresponding to a 470-μm displacement at the stimulation point in the previously calibrated actuator input–output relationship.[2] The same stimulus intensity was applied across all postoperative time points and experimental groups to provide a standardized assay of TC potentiation. Mice were anesthetized with urethane (1.25 g/kg, intraperitoneal; Sigma-Aldrich), mounted in a stereotaxic apparatus (51730; Stoelting), and maintained at 37 °C. Signals were acquired at 30 kHz using a SmartBox Pro data-acquisition system and Allego software (NeuroNexus Technologies). Probe impedance (1–2.5 MΩ) was measured in PBS before recording as described previously.[2] Probes were coated with the lipophilic carbocyanine dye 1,1′-dioctadecyl-3,3,3′,3′-tetramethylindocarbocyanine perchlorate (DiI) for *post hoc* localization. Ag/AgCl pellets (EP1; World Precision Instruments) positioned distal to the recording site served as ground and reference electrodes. In simultaneous VPM–L4 experiments, one probe was targeted to barrel cortex (AP, −1.5 mm; ML, 3.2 mm; DV, −1.25 mm; 30° insertion angle) and a second probe to VPM (AP, −1.8 mm; ML, 1.45 mm; DV, −4.0 mm) in the same animal.

For Neuropixels recordings, extracellular activity was acquired from barrel cortex and VPM using Neuropixels 1.0 probes[7] connected to a Neuropixels headstage and PXIe acquisition module (PXIe_1000; imec) through the Neuropixels interface cable. Barrel cortex recordings were targeted to wS1 (AP, −1.7 mm; ML, 3.2 mm; DV, 3.5 mm; 30° insertion angle), whereas VPM recordings were targeted using stereotaxic and histological landmarks. Recordings were controlled using SpikeGLX. Neural activity was recorded during a behavioral threshold-testing session comprising catch trials and pseudorandomly interleaved whisker deflections of 0.3°, 0.4°, 0.5°, 0.6°, 0.7°, 0.8°, 0.9°, 1.0°, 1.3°, 1.8°, 3.3°, 5.5°, 8.3°, and 11°. Each of the 15 conditions, comprising catch trials and 14 nonzero stimulus intensities, was presented 20 times, yielding 300 trials in total. The 2.5° stimulus was used during sequential behavioral training but was not included as a separate Neuropixels test intensity. Trial events were aligned to the digital rising edge marking whisker-stimulus onset.

Neuropixels recordings were performed after mice reached learning criterion, between PO33 and PO70, a period selected to assess the sustained phase of learning-associated TC potentiation in IONX mice.

**Spike sorting and spike-count analysis**

Raw data were processed offline with Kilosort4[8] followed by manual or semi-automated curation with Phy. Well-isolated units labeled "good" were used as the primary single-unit dataset. Multi-unit activity was included for population-level recruitment analyses, as specified in the figure legends. Units failing quality-control criteria were excluded. Sorted data were analyzed using custom Python scripts. Peri-stimulus analyses used −200 ms to +1,000 ms relative to stimulus onset. Stimulus-evoked L4 spike counts were computed across whisker deflection amplitudes. Recruitment probability was quantified as the proportion of trials containing an above-baseline response, defined unless otherwise stated as trials in which post-stimulus spike count exceeded baseline spike count within the same trial. Conditional response magnitude was assessed using response-positive trials only. Sensory responses of wS1 and VPM were quantified in the 0–50 ms and 0–20 ms windows, respectively. Baseline spike counts were measured during the pre-stimulus period and duration-matched to the corresponding evoked window. ΔL4 spikes per trial was calculated as the 0–50-ms post-stimulus spike count minus the duration-matched baseline count; ΔVPM spikes per trial was calculated analogously using the 0–20-ms post-stimulus window. When baseline counts were available only for the −50 to 0 ms or −200 to 0 ms pre-stimulus interval, they were scaled to the duration of the corresponding evoked window. For Neuropixels population analyses, single-unit and multi-unit responses were treated as observations nested within mice. To avoid pseudoreplication, stimulus-evoked L4 responses for Fig. 5g and related stimulus-response analyses were averaged across validated L4 single-unit and multi-unit activity sites within each mouse at each stimulus intensity before group-level statistical testing. Statistical inference was performed on these mouse-level values, with mouse identity defining the biological replicate.

**Local field potential analysis**

For local field potential (LFP) analysis, raw signals were downsampled to 1 kHz and band-pass filtered between 0.1 and 300 Hz. Stimulus-aligned LFP traces were baseline-corrected using the mean voltage during the −10 to 0 ms pre-stimulus period and averaged across trials.

Whisker-evoked peak amplitude was defined as the magnitude of the largest negative-going deflection within 0–50 ms after stimulus onset in L4 and within 0–20 ms after stimulus onset in VPM. Because the evoked LFP response was negative-going, the minimum voltage was multiplied by −1 and reported as a positive peak amplitude in microvolts. Peak amplitudes were averaged across trials within each mouse and experimental condition.

**Laminar localization and current-source-density analysis**

Layer assignments were anchored using probe depth and current-source density (CSD) organization. Stimulus-aligned LFP epochs were baseline-corrected using the −10 to 0 ms pre-stimulus window and averaged across trials. For Neuropixels CSD visualization, channels at the same vertical depth were averaged across probe columns, and the resulting depth-by-time LFP matrix was smoothed across depth and time before CSD calculation. CSD was calculated as

$$CSD(r,t) = -[\Phi(r+h,t) - 2\Phi(r,t) + \Phi(r-h,t)] / h^2,$$

where $\Phi(r,t)$ is the LFP at depth r and time t, and h is the median inter-depth spacing after depth alignment. With this sign convention, positive CSD values were plotted and interpreted as sinks and negative values as sources. The principal early whisker-evoked sink was used as the physiological landmark for the thalamocortical input layer and validated L4 localization. Neuropixels L4 units were assigned using probe depth relative to this physiological anchor, supported by histology or depth mapping when available.

**Threshold analyses**

Perceptual thresholds during Neuropixels testing were estimated from the relationship between behavioral performance and stimulus-evoked L4 population recruitment across whisker amplitudes. Behavioral performance was quantified as lick-response probability at each tested intensity, with catch trials included as the 0° condition. For each mouse and stimulus condition, one neural–behavioral point was defined by the mouse-level mean stimulus-evoked L4 Δ spike count per trial in the 0–50-ms post-stimulus window and the corresponding lick-response probability.

For each mouse, stimulus conditions were ordered by increasing intensity. Neural and behavioral coordinates were independently range-scaled within mouse so that neither axis dominated the fit. Each admissible internal transition point was evaluated with a two-segment total least-squares model that minimized the summed orthogonal residual error across the two trajectory segments. The tested intensity corresponding to the minimum-error transition was defined as the total least-squares-derived perceptual threshold, provided that the post-transition segment showed a steeper behavioral increase than the pre-transition segment. Group differences were assessed using a two-sided exact permutation test based on the Mann–Whitney U statistic, with mouse identity defining the biological replicate. As a sensitivity analysis, transition points were also assigned by visual inspection using the same qualitative transition criterion. Manual and total least-squares-derived estimates differed by ≤0.2° in 14 of 15 mice. Segmented total least-squares fitting was restricted to tested intensities ≤1.8° to focus the analysis on the low-intensity transition region and avoid higher-intensity, near-asymptotic responses from dominating the fit.

**Temporal sharpening analysis**

Temporal sharpening index (TSI) was calculated as the number of sensory-evoked spikes in the peak 10-ms window within the 0–100-ms post-stimulus interval divided by the total number of sensory-evoked spikes in that interval. This analysis was motivated by prior work showing that millisecond-scale spike timing and temporal response patterns can contribute to sensory coding.[9,10] For each unit and stimulus intensity, the peak 10-ms window was identified by sliding a 10-ms window across the 0–100-ms interval and selecting the window containing the maximum sensory-evoked spike count. Unit-level TSI values were then averaged within each mouse at each stimulus intensity. ΔTSI was calculated for each mouse as mean TSI at high stimulus intensities, defined as 3.3°, 5.5°, 8.3°, and 11°, minus mean TSI at low stimulus intensities, defined as 0.3°, 0.4°, and 0.5°. Group-level comparisons were performed on these mouse-level ΔTSI values, so that the animal, rather than the number of recorded units, defined the independent biological replicate. Spike trains were binned at 5 ms before scanning with the 10-ms sliding window.

**Histology and anatomical verification**

Recording locations, probe trajectories, drug-delivery sites, and laminar landmarks were verified histologically after perfusion when applicable. Histology was used to confirm barrel cortex targeting, VPM localization, laminar positioning, and pump or probe placement.

VPM barreloid and wS1 barrel-field mapping

Cytochrome oxidase histochemistry, fluorescence imaging, and DiI probe-track labeling were used to identify VPM barreloids, wS1 barrels, and electrophysiological recording sites. For VPM barreloid mapping, fixed brains were sectioned in an oblique plane selected to expose the VPM barreloid array, guided by prior anatomical descriptions of barreloid organization in the rodent thalamus.[11] Sections were processed for cytochrome oxidase activity, mounted, and imaged to identify the barreloid field. After the cytochrome oxidase-defined barreloid region was identified, the corresponding region was examined at higher magnification using green autofluorescence to visualize the barreloid pattern.

**Anterograde VPM axon tracing and DiI probe-track validation**

To identify the thalamocortical projection zone in wS1, an anterograde adeno-associated virus serotype 9 (AAV9) vector expressing mCherry (AAV9.CaMKIIa.hChR2(E123A).mCherry.WPRE.hGH; RRID: Addgene_35506)[12] was injected into VPM. Mice were anesthetized with 2–3% isoflurane in oxygen and placed in a stereotaxic frame. A small craniotomy was made above VPM, and virus was delivered using a 10-μL glass syringe fitted with a 33-gauge injection needle. The injection was targeted to VPM using stereotaxic coordinates approximately 1.8 mm posterior and 1.4 mm lateral to bregma, with the needle lowered 3.5 mm below the brain surface. A total volume of 100 nL virus was injected. The injection needle was left in place for 10 min before slow withdrawal to minimize backflow. The craniotomy was sealed with bone wax, the scalp was sutured, and mice were allowed to recover.

Virus expression proceeded for approximately 2 weeks before histological analysis. Mice were deeply anesthetized and transcardially perfused with PBS followed by fixative. Brains were post-fixed overnight, and coronal sections were cut through wS1 and VPM using a vibratome. Sections were mounted, coverslipped with antifade mounting medium containing 4′,6-diamidino-2-phenylindole (DAPI), and imaged by fluorescence microscopy. mCherry-positive VPM axons were used to delineate the thalamocortical projection zone in wS1. Dense, drop-like VPM axon labeling in L4, together with deeper labeling in L5, was used as an anatomical landmark for the wS1 barrel field. Recording sites were verified by coating probes with DiI before insertion and imaging the resulting fluorescent probe tracks after fixation. wS1 recordings were included when the DiI-labeled trajectory intersected the barrel cortex recording region, and VPM recordings were included when the probe track passed through the VPM barreloid region.

**Cytochrome oxidase staining for wS1 L4 barrel localization**

Cytochrome oxidase histochemistry was used to visualize the wS1 barrel field and confirm the tangential location of Neuropixels probe tracks within L4. After completion of Neuropixels recordings, mice were deeply anesthetized and transcardially perfused with PBS followed by fixative. The cortex containing wS1 was dissected, flattened, and post-fixed, and tangential sections through L4 were cut on a vibratome. Sections were processed for cytochrome oxidase activity using a reaction solution containing cytochrome c, 3,3′-diaminobenzidine, and catalase in phosphate buffer. Stained sections were mounted and imaged to identify the barrel-row and barrel-column location of the probe track relative to the wS1 barrel map. Recordings were included when histology and physiological depth mapping confirmed probe placement within the targeted wS1 L4 barrel region.

**Randomization, blinding, and exclusion criteria**

Animals were assigned to experimental groups while balancing litter, sex, and surgery date. Investigators were necessarily aware of group assignment during Sham or IONX surgery because the procedures differed. Behavioral, electrophysiological, and histological analyses used predefined criteria, with coded animal identifiers applied when technically feasible. Animals or sessions were excluded only for technical reasons, including unsuccessful recovery, recording or pump failure, unconfirmed recording location, or incomplete data caused by technical interruption.

**Statistics and reproducibility**

Statistical analyses were performed using Prism, R, and Python. Two-group comparisons used parametric or nonparametric tests depending on data structure and distribution. For Supplementary Fig. 4d–f,h, group differences in response amplitudes and VPM–L4 input–output slopes were assessed using two-sided Welch's two-sample t-tests because sample sizes were unequal and variance estimates differed between groups. Mouse-level values were used as the unit of inference, and nonparametric sensitivity analyses were reported in the Source Data where applicable. Multi-group comparisons used ANOVA or nonparametric alternatives with multiple-comparison correction where appropriate. Repeated behavioral trajectories were analyzed using repeated-measures, mixed-effects, or GEE models. Learning-completer proportions were assessed using Fisher's exact test. Time-to-criterion analyses used log-rank Mantel–Cox tests when non-learners were censored. HMM-derived learning onset, learning completion, and consolidation interval were assessed using two-sided nonparametric tests when distributions were skewed. Psychometric and stimulus-response curves were analyzed using mixed ANOVA or mixed-effects models with group as a between-subject factor and stimulus intensity as a repeated factor, followed by Holm-corrected *post hoc* comparisons across stimulus intensities. For datasets containing repeated observations within animals, including stimulus intensities, learning states, retention time points, and Neuropixels recordings, mouse identity defined the biological replicate. Neuropixels stimulus-response statistics were performed on mouse-level values after averaging validated L4 single-unit and multi-unit activity responses within each mouse at each stimulus intensity. Unless otherwise stated, summary data are presented as mean ± SEM. Exact P values, sample sizes, effect estimates, and confidence intervals, where applicable, are reported in the figure legends and Source Data. No statistical method was used to predetermine sample size; sample sizes were chosen on the basis of prior studies using similar designs and are reported for each experiment.

### Supplementary Figures

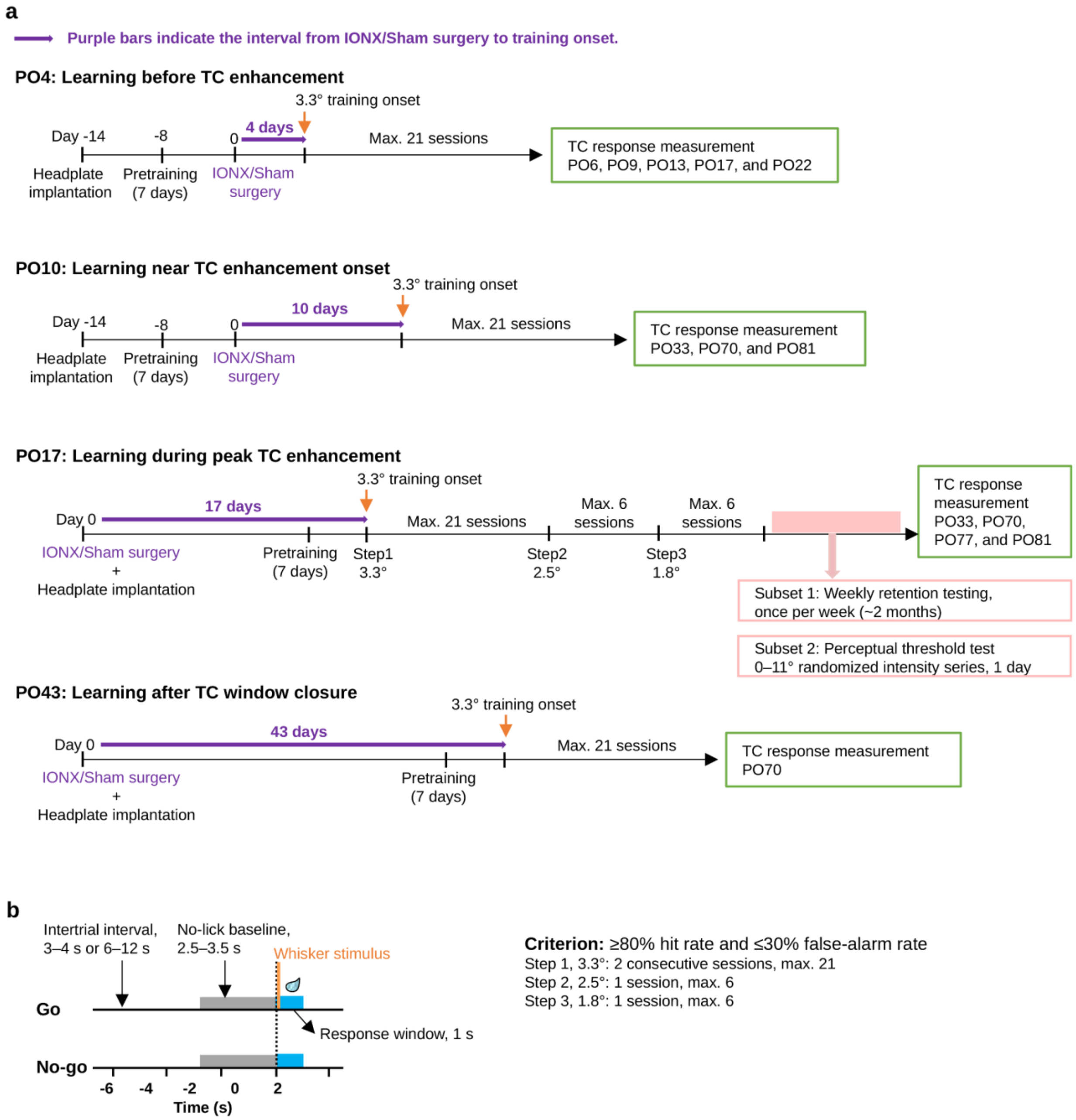


**Supplementary Fig. 1 | Experimental design for learning-timing experiments. Related to Figs. 1 and 6.**

(a) Experimental timelines for 3.3° whisker detection training initiated at different postoperative phases after IONX or Sham surgery. Training began at PO4, before detectable TC enhancement;

PO10, near the onset of TC enhancement; PO17, during peak TC enhancement; or PO43, after closure of the TC enhancement window. Purple bars indicate the interval from IONX or Sham surgery to training onset. Green boxes indicate postoperative time points used for whisker-evoked TC response measurements. PO17-trained mice were assigned to weekly retention testing or perceptual threshold testing after sequential training at lower intensities.

(b) Go/No-go trial structure and learning criteria. On Go trials, a whisker stimulus was delivered after a no-lick baseline period, and licking during the 1-s response window was scored as a hit. On No-go trials, no stimulus was delivered, and licking during the corresponding response window was scored as a false alarm. The intertrial interval was initially 3–4 s and was increased to 6–12 s when impulsive licking increased. Learning criterion was ≥80% hit rate and ≤30% false-alarm rate for two consecutive sessions at 3.3° and for one session at 2.5° and 1.8°, with training limited to six sessions at each lower intensity.

Abbreviations: IONX, infraorbital nerve transection; TC, thalamocortical; PO, postoperative day.

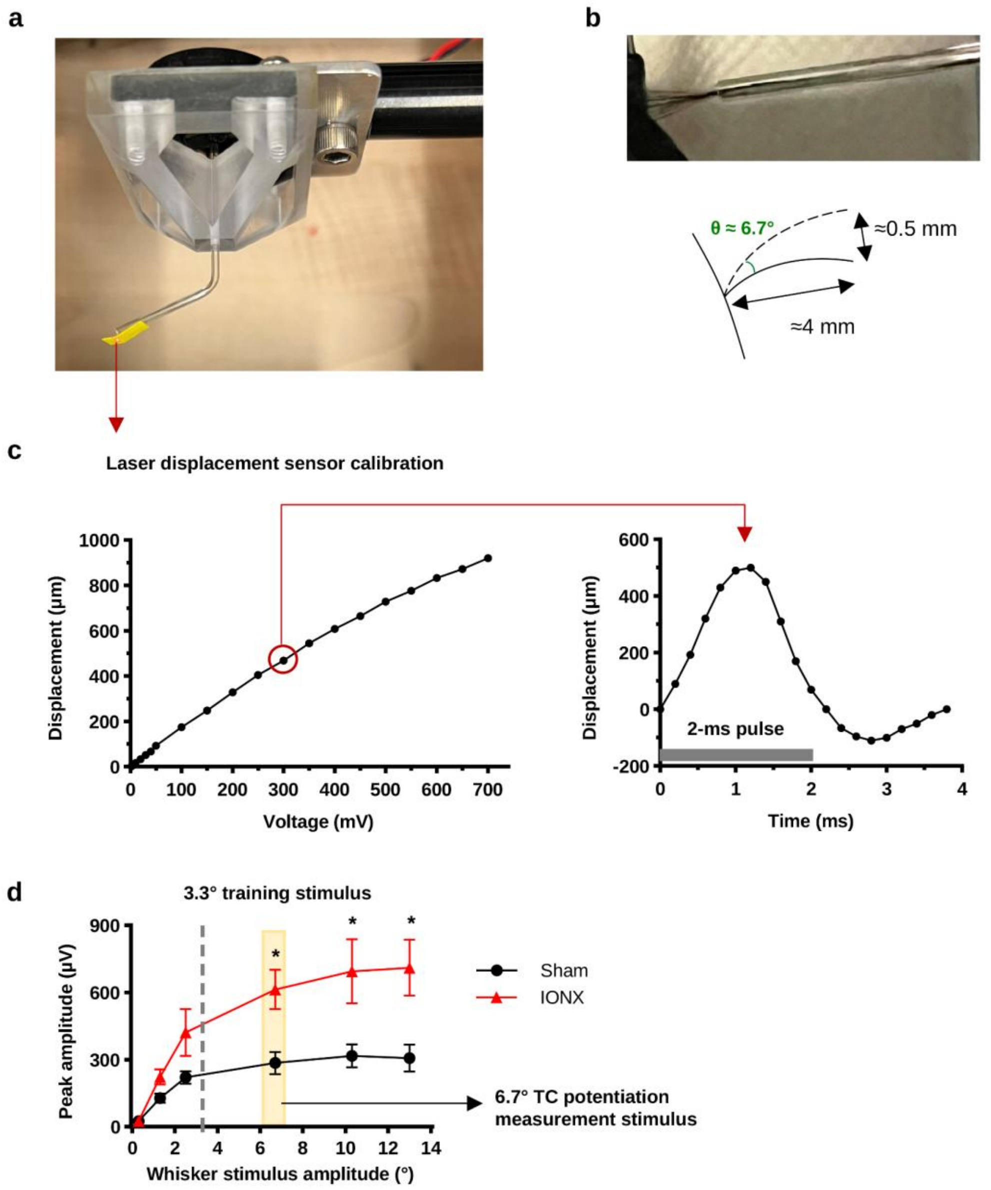


**Supplementary Fig. 2 | Laser calibration of the coil-driven whisker actuator and stimulus selection. Related to Fig. 1.**

(a) Custom coil-driven actuator used to deliver brief whisker deflections during head-fixed behavior. A capillary glass tube was attached to the voice coil for whisker insertion.

(b) Whisker insertion and angular-deflection geometry. Selected whiskers were inserted into the capillary tube approximately 4 mm from the whisker pad, and angular deflection was calculated from measured actuator displacement and stimulation distance.

(c) Laser displacement calibration of the actuator performed for the present study. Left, command voltage–displacement relationship measured with a laser displacement sensor; right, representative actuator displacement trace evoked by a 2-ms command pulse.

(d) Whisker stimulus input–output relationship modified from Supplementary Ref. 2 and replotted after conversion of displacement to angular deflection using the present actuator geometry. The 6.7° stimulus (470-μm displacement in Supplementary Ref. 2) was used for TC potentiation measurements because it produced a stable, high-amplitude L4 LFP response and robust IONX/Sham separation. The 3.3° stimulus was used for behavioral training because it fell within a non-saturating range suitable for detecting IONX-dependent learning advantages. Data are mean ± SEM (Sham, n = 5 mice; IONX, n = 6 mice). Group differences were assessed using two-sided Mann–Whitney U-tests with Holm correction across stimulus amplitudes (6.7°, adjusted $P = 0.04329$; 10.3°, adjusted $P = 0.04329$; 13.0°, adjusted $P = 0.02597$). *$P < 0.05$. Abbreviations: IONX, infraorbital nerve transection; TC, thalamocortical; L4, layer 4; LFP, local field potential.

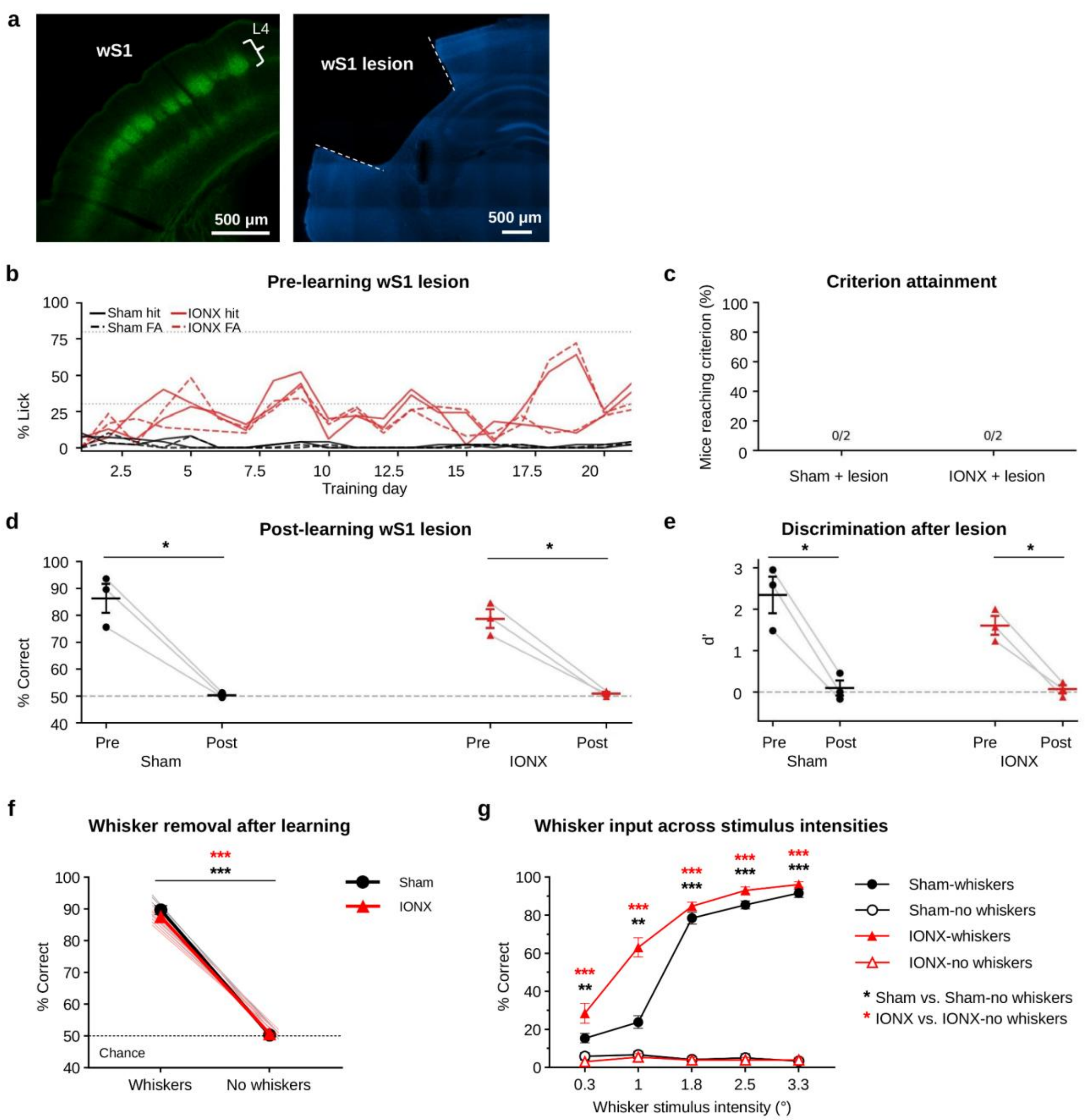


**Supplementary Fig. 3 | Learned whisker detection requires whisker-mediated tactile input and contralateral wS1. Related to Fig. 1.**

(a) Representative histological images showing intact wS1 and wS1 lesion. Dashed outlines indicate the wS1 region. Scale bars, 500 µm.

(b) Lick probability across training in mice with wS1 lesion before learning. Solid lines indicate hit rate on Go trials, and dashed lines indicate false-alarm rate on No-go trials.

(c) Criterion attainment after pre-learning wS1 lesion. No Sham + lesion or IONX + lesion mice reached criterion during 3.3° whisker-detection training (Sham + lesion, 0 of 2 mice; IONX + lesion, 0 of 2 mice; two-sided Fisher's exact test, $P = 1.0$).

(d) Behavioral performance before and after post-learning wS1 lesion in mice that had reached criterion (Sham, $n = 3$ mice; IONX, $n = 3$ mice). Lines connect measurements from the same mouse. wS1 lesion reduced performance to chance in both groups (paired two-sided t-tests: Sham, $P = 0.01899$; IONX, $P = 0.01324$).

(e) Discrimination performance before and after post-learning wS1 lesion in the same mice. Lines connect measurements from the same mouse. wS1 lesion reduced d′ to near zero in both groups (paired two-sided t-tests: Sham, $P = 0.01696$; IONX, $P = 0.01544$).

(f) Behavioral performance before and after trained-whisker removal in the 8.3° cohort (Sham, $n = 7$ mice; IONX, $n = 8$ mice). Lines connect measurements from the same mouse. Whisker removal abolished task performance in both groups (paired two-sided t-tests: Sham, $P = 1.48 \times 10^{-7}$; IONX, $P = 5.27 \times 10^{-11}$).

(g) Detection performance across whisker stimulus intensities before and after removal of the trained whiskers in the 3.3° cohort (Sham, $n = 12$ mice; IONX, $n = 13$ mice). Filled symbols indicate whiskers-intact testing, and open symbols indicate no-whisker testing. Within each group, whiskers-intact and no-whisker performance was compared at each of the five stimulus intensities using paired two-sided t-tests. P values were Holm-adjusted separately across the five intensity-wise comparisons within the Sham and IONX groups.

Data are mean ± SEM where summary values are shown. Individual lines or points represent mice. *$P < 0.05$, **$P < 0.01$, ***$P < 0.001$. Abbreviations: wS1, whisker primary somatosensory cortex; IONX, infraorbital nerve transection.

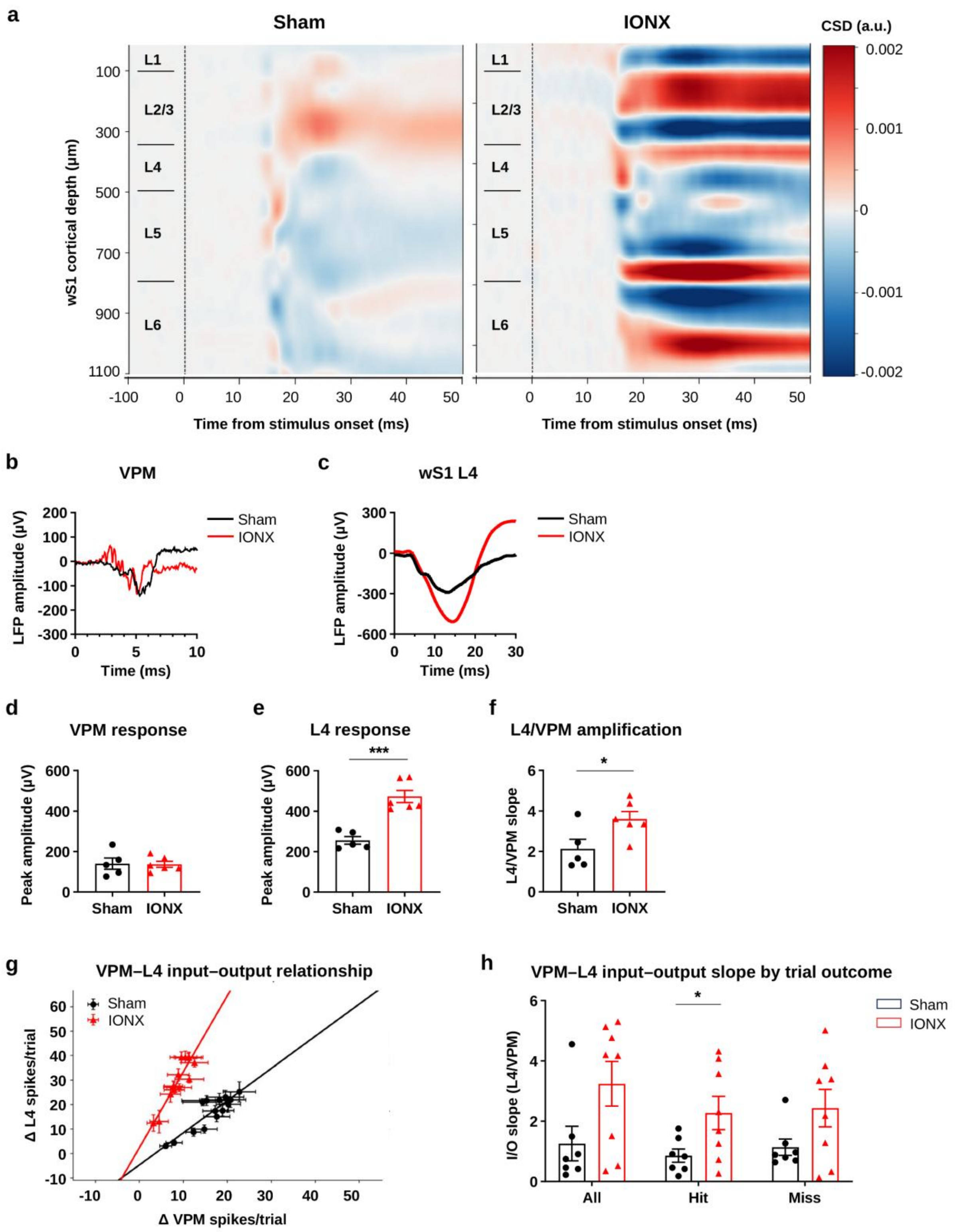


**Supplementary Fig. 4 | IONX enhances cortical amplification of VPM input. Related to Figs. 1, 4, and 5.**

(a) Whisker stimulus-aligned current-source density (CSD) maps used to identify the principal early L4 thalamocortical input sink and assign recording depths. Positive CSD values denote sinks and negative values denote sources.

(b) Representative whisker-evoked VPM local field potential responses.

(c) Representative whisker-evoked L4 local field potential responses.

(d) VPM response amplitude (Sham: 140.53 ± 27.69, n = 5 mice; IONX: 137.30 ± 14.78, n = 6 mice; two-sided Welch's t-test, t = 0.103, P = 0.921).

(e) L4 response amplitude. IONX elevated responses (Sham: 256.08 ± 19.22, n = 5 mice; IONX: 473.08 ± 30.07, n = 6 mice; Welch's t-test, t = −6.08, $P = 2.67 \times 10^{-4}$).

(f) L4/VPM amplification slope (Sham: 2.12 ± 0.48, n = 5 mice; IONX: 3.61 ± 0.36, n = 6 mice; Welch's t-test, t = −2.48, P = 0.0386).

(g) VPM–L4 input–output relationship across intensities using stimulus-evoked spike counts quantified in region-specific post-stimulus windows (VPM, 0–20 ms; L4, 0–50 ms; lines indicate linear fits per mouse).

(h) VPM–L4 input–output slopes across trial types. IONX increased the slope during hit trials (Sham: 0.86 ± 0.22, n = 7 mice; IONX: 2.28 ± 0.55, n = 8 mice; Welch's t-test, t = −2.39, P = 0.0400). Slope did not significantly differ across all trials (P = 0.0544) or miss trials (P = 0.0847).

Data are mean ± SEM. Points in d–f and h represent mice. *P < 0.05; ***P < 0.001.

Abbreviations: IONX, infraorbital nerve transection; VPM, ventral posteromedial thalamic nucleus; L4, layer 4; CSD, current-source density; TC, thalamocortical.

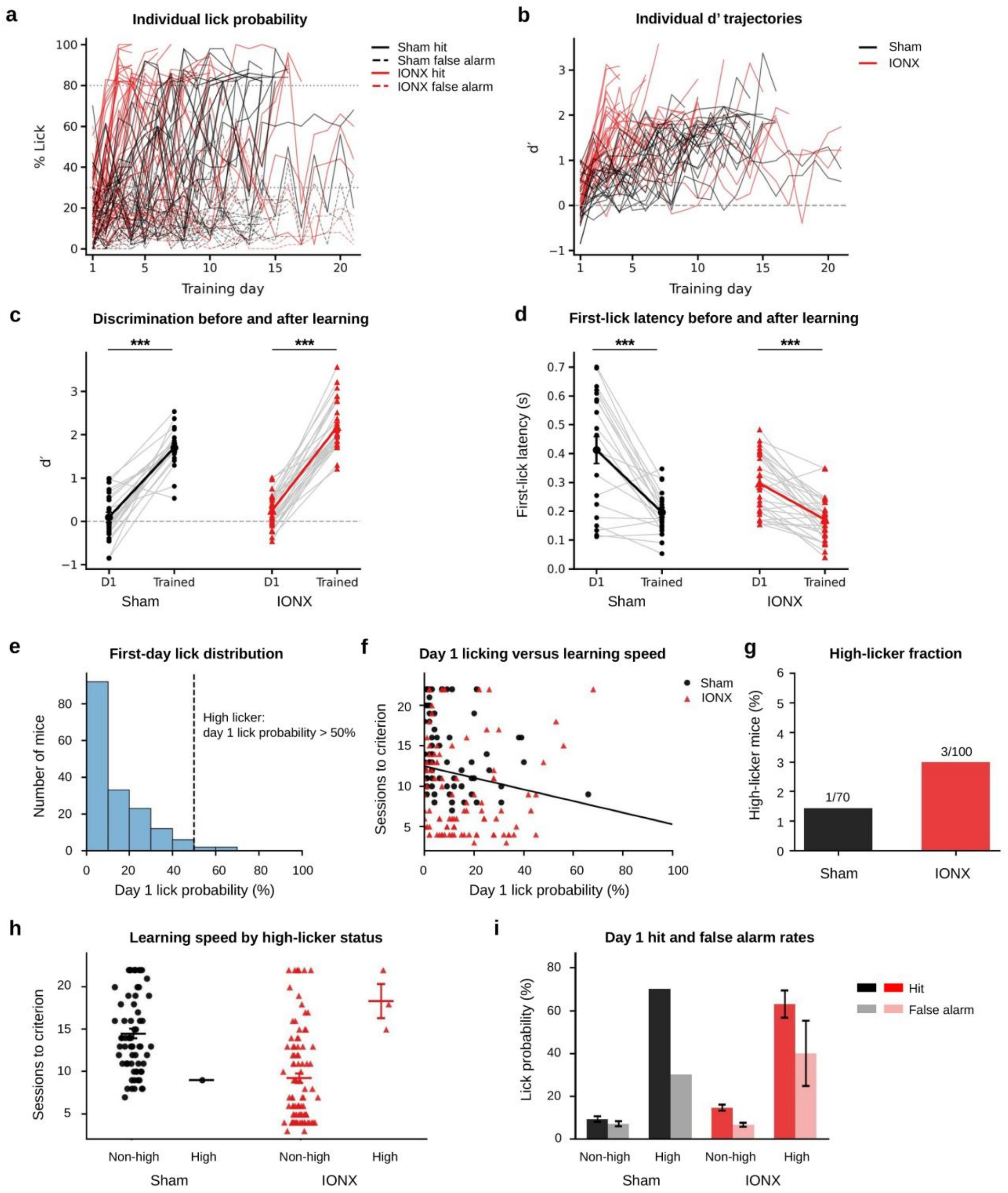


**Supplementary Fig. 5 | Behavioral controls for accelerated whisker detection learning after IONX. Related to Figs. 1 and 2.**

(a) Lick probability across 3.3° training at PO17 (solid: hit rate; dashed: false-alarm rate; horizontal lines: criteria).

(b) Individual d′ trajectories (dashed line: d′ = 0).

(c) Discrimination (d′) on Day 1 (D1) versus the trained state (Sham, n = 22 mice; IONX, n = 29 mice). Performance increased in both groups (paired two-sided t-tests: Sham, $P = 1.11 \times 10^{-9}$; IONX, $P = 4.40 \times 10^{-17}$).

(d) First-lick latency on D1 versus the trained state (Sham, n = 20 mice; IONX, n = 29 mice). Latency decreased significantly after learning in both groups (paired two-sided t-tests: Sham, $P = 4.08 \times 10^{-5}$; IONX, $P = 2.15 \times 10^{-7}$).

(e) Distribution of D1 lick probability across cohorts (dashed line: 50% high-licker cutoff).

(f) D1 lick probability versus sessions-to-criterion plotting value (descriptive linear regression; non-learners assigned 22 sessions; $R^2 = 0.030$, $P = 0.0240$).

(g) High-licker fraction (Sham: 1 of 70 mice; IONX: 3 of 100 mice; two-sided Fisher's exact test, $P = 0.644$).

(h) Sessions to criterion stratified by high-licker status. The IONX–Sham difference remained significant after excluding high-lickers (Sham, n = 69 mice; IONX, n = 97 mice; two-sided Mann–Whitney U-test, U = 5,183, $P = 1.60 \times 10^{-9}$), whereas pooled high- and non-high-lickers did not differ significantly (n = 4 and n = 166 mice, respectively; U = 181.5, $P = 0.122$).

(i) D1 hit and false-alarm rates in non-high-lickers and high-lickers. No inferential statistical test was performed for this descriptive analysis.

Data are mean ± SEM. Lines/points represent individual mice. ***$P < 0.001$. Abbreviations: IONX, infraorbital nerve transection; PO, postoperative day; d′, sensitivity index.

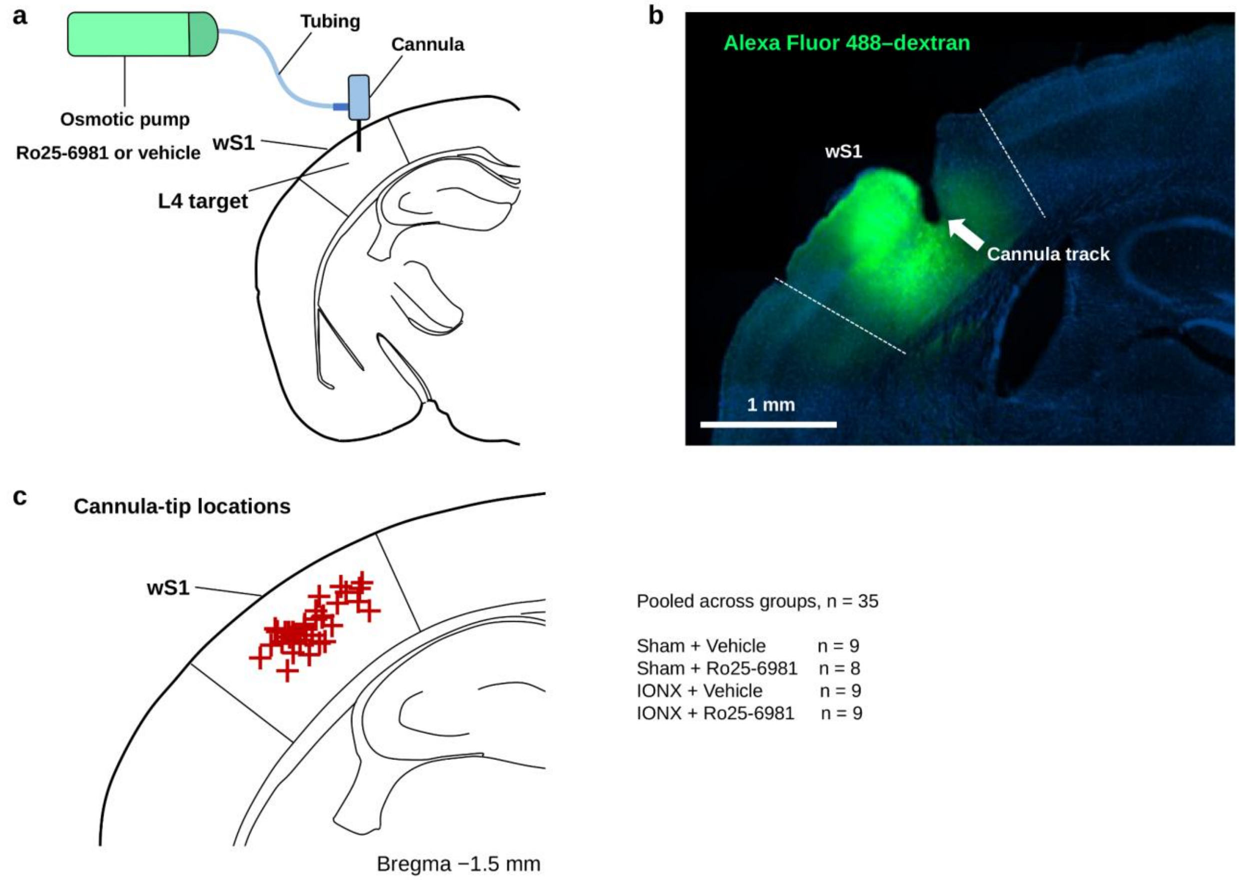


**Supplementary Fig. 6 | Histological assessment of local infusion targeting in wS1. Related to Fig. 3.**

(a) Schematic of osmotic pump infusion into wS1. Ro25-6981 or vehicle was delivered through an implanted cannula targeting L4 of wS1. Alexa Fluor 488–dextran was infused using the same pump and cannula configuration to estimate local infusion spread.

(b) Representative coronal section from a separate validation animal showing Alexa Fluor 488–dextran spread after osmotic pump delivery into wS1. The cannula track and dextran fluorescence overlapped the intended dorsolateral wS1 target region. Scale bar, 1 mm.

(c) Histologically verified cannula-tip locations for all behavioral pharmacology animals (n = 35 mice total: Sham + Vehicle, n = 9; Sham + Ro25-6981, n = 8; IONX + Vehicle, n = 9; IONX + Ro25-6981, n = 9). Each cross represents one mouse.

Abbreviations: wS1, whisker primary somatosensory cortex; L4, layer 4; IONX, infraorbital nerve transection.

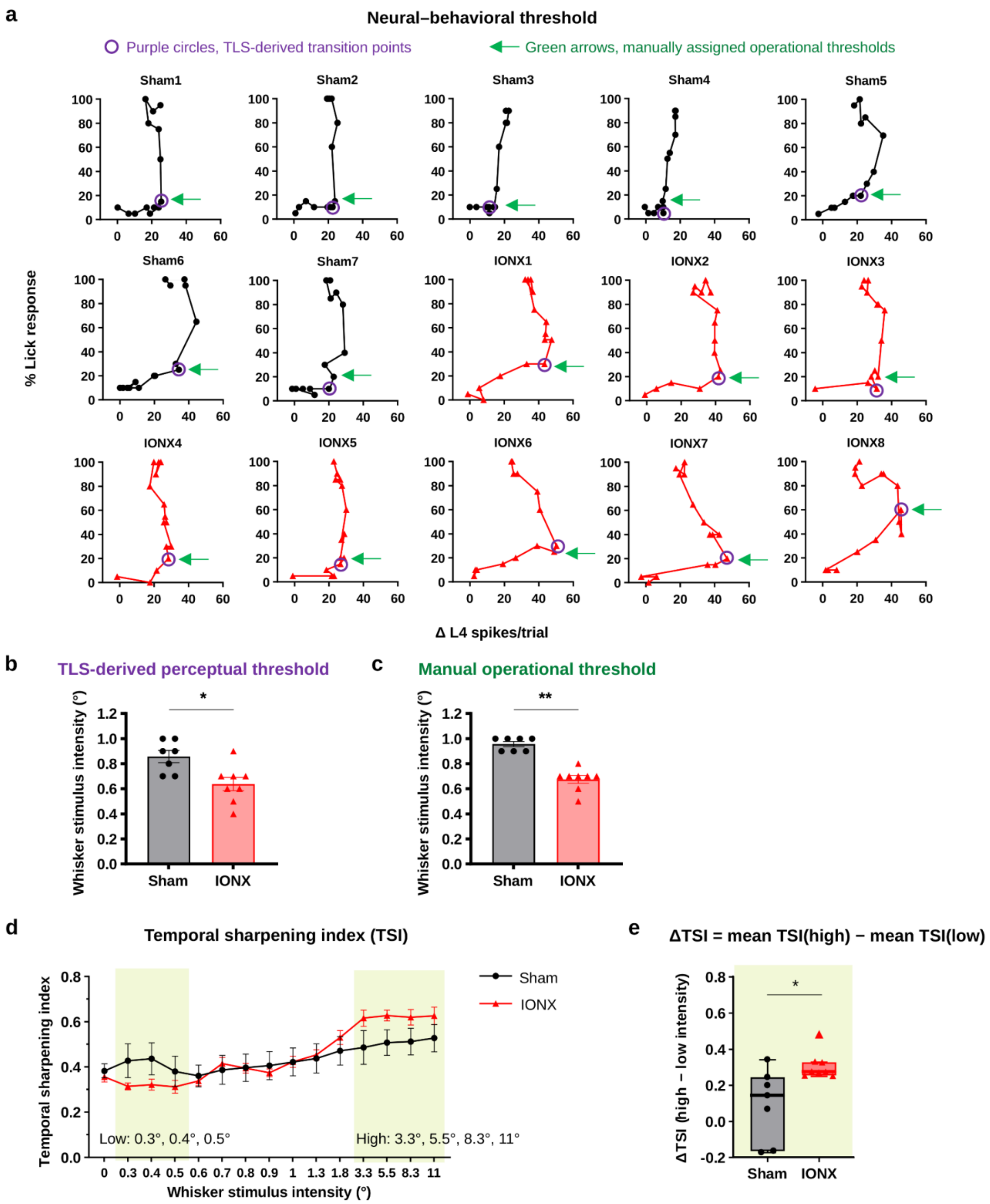


**Supplementary Fig. 7 | Neural–behavioral threshold and temporal sharpening analyses.**

**Related to Fig. 5.**

(a) Individual neural–behavioral trajectories from Sham and IONX mice. Each panel plots lick-response probability against stimulus-evoked L4 Δ spikes per trial for one mouse, with points connected in order of increasing stimulus intensity. Purple circles indicate segmented total least-squares-derived transition points, and green arrows indicate manually assigned operational thresholds. The two estimates differed by ≤0.2° in 14 of 15 mice.

(b) Total least-squares-derived perceptual thresholds were lower after IONX (Sham, 0.857 ± 0.048°, n = 7 mice; IONX, 0.638 ± 0.053°, n = 8 mice; two-sided exact permutation test based on the Mann–Whitney U statistic, U = 49.0, P = 0.0121).

(c) Manually assigned operational thresholds were also lower after IONX (Sham, 0.957 ± 0.020°, n = 7 mice; IONX, 0.675 ± 0.031°, n = 8 mice; two-sided Mann–Whitney U-test, U = 56.0, P = 0.00102).

(d) Temporal sharpening index (TSI) across whisker stimulus intensities. TSI was calculated as the peak 10-ms spike count divided by total spike count in the 0–100-ms post-stimulus window.

(e) Mouse-level ΔTSI, calculated as mean TSI at high stimulus intensities minus mean TSI at low stimulus intensities. Low intensities were 0.3°, 0.4°, and 0.5°; high intensities were 3.3°, 5.5°, 8.3°, and 11°. IONX increased ΔTSI relative to Sham controls (Sham, 0.094 ± 0.075, n = 7 mice; IONX, 0.306 ± 0.027, n = 8 mice; two-sided Mann–Whitney U-test, U = 7.0, P = 0.01399).

Abbreviations: IONX, infraorbital nerve transection; L4, layer 4; TSI, temporal sharpening index.

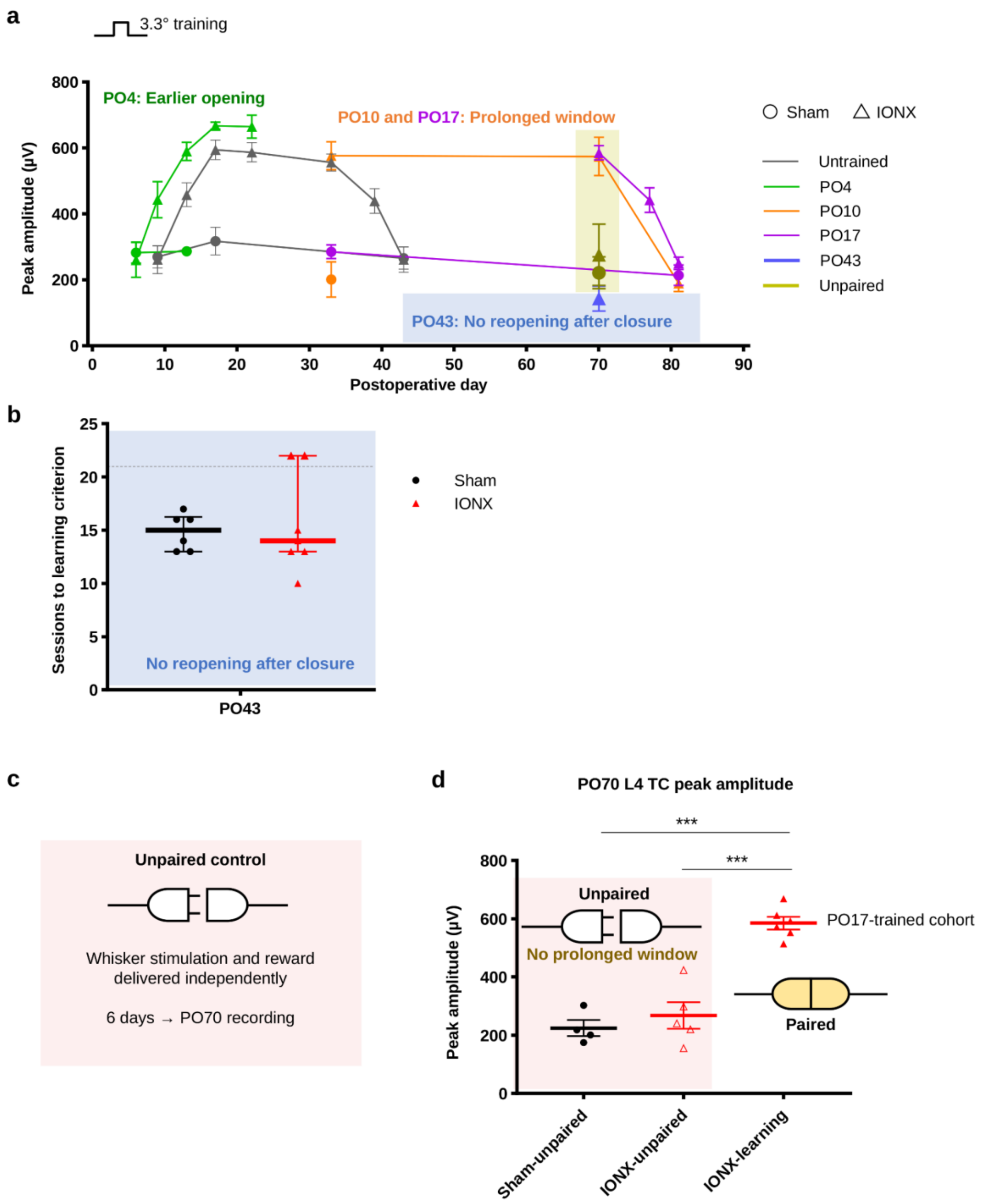


**Supplementary Fig. 8 | Learning timing and sensory–reward contingency shape the reactivated thalamocortical plasticity window. Related to Fig. 6.**

(a) Summary of whisker-evoked L4 TC response amplitude across postoperative time under different learning-timing conditions. Training initiated at PO4 advanced the emergence of TC enhancement, whereas training initiated at PO10 or PO17 prolonged it. Training initiated at PO43 did not reopen the window after closure.

(b) Criterion attainment after training initiated at PO43, after closure of the TC enhancement window. Six of six Sham mice and five of seven IONX mice reached criterion (two-sided Fisher's exact test, $P = 0.4615$). Criterion-attainment timing did not differ between groups (Sham, $n = 6$ mice; IONX, $n = 7$ mice; log-rank Mantel–Cox test, $\chi^2 = 0.1141$, d.f. = 1, $P = 0.736$). Mice that did not reach criterion within the 21-session training period were treated as right-censored; values of 22 sessions are shown only for visualization.

(c) Unpaired control design. Whisker stimulation and reward were delivered independently for 6 days before PO70 recording.

(d) PO70 L4 TC peak amplitude after paired learning or unpaired sensory and reward exposure (Sham-unpaired, $n = 4$ mice; IONX-unpaired, $n = 5$ mice; IONX-learning, $n = 6$ mice). Groups differed significantly by one-way ANOVA ($F(2,12) = 38.55$, $P = 5.97 \times 10^{-6}$). Tukey's multiple-comparison test showed that IONX-learning responses were higher than those in both Sham-unpaired and IONX-unpaired mice (both adjusted $P < 0.0001$), whereas Sham-unpaired and IONX-unpaired groups did not differ (adjusted $P = 0.6578$). Thus, unpaired sensory stimulation and reward exposure did not maintain the prolonged TC state.

Abbreviations: IONX, infraorbital nerve transection; L4, layer 4; TC, thalamocortical; PO, postoperative day.

**Supplementary Table 1 | Key resources**

| **REAGENT or RESOURCE** | **SOURCE** | **IDENTIFIER** |
|---|---|---|
| **Bacterial and virus strains** | | |
| pAAV-CaMKIIa-hChR2(E123A)-mCherry (AAV9) | Addgene | Addgene viral prep #35506-AAV9; RRID:Addgene_35506 |
| **Chemicals, peptides, and recombinant proteins** | | |
| Ro25-6981 maleate | Tocris Bioscience | Cat# 1594 |
| Dextran, Alexa Fluor 488, molecular weight 10,000, anionic, fixable | Invitrogen, Thermo Fisher Scientific | Cat# D22910 |
| Urethane, ≥99% | Sigma-Aldrich | Cat# U2500 |
| C&B Metabond Quick Adhesive Cement System | Parkell | SKU S380 |
| Cytochrome c from equine heart, ≥95% (SDS-PAGE) | Sigma-Aldrich | Cat# C2506 |
| 3,3′-Diaminobenzidine tetrahydrochloride hydrate, ≥96% | Sigma-Aldrich | Cat# D5637 |
| Catalase from bovine liver, lyophilized powder | Sigma-Aldrich | Cat# C9322 |
| Deposited data | | |

| REAGENT or RESOURCE | SOURCE | IDENTIFIER |
|---|---|---|
| Source data underlying figures | This paper | Provided with this paper |
| **Experimental models: Organisms/strains** | | |
| Mouse: C57BL/6J, both sexes | The Jackson Laboratory; bred in-house | Stock No. 000664; RRID:IMSR_JAX:000664 |
| **Software and algorithms** | | |
| LabVIEW 2019 SP1 | National Instruments | Version 19.0.1f5 |
| SOLIDWORKS 3D CAD software | Dassault Systèmes SolidWorks Corp. | Version 2024 SP5.0 |
| SpikeGLX | Janelia Research Campus | Release 20240620-phase30 |
| imec API | imec | Version 3.70.2 |
| Allego | NeuroNexus Technologies | Version 3.4.4 |
| Kilosort4 | Pachitariu et al.[8] | Version 4.0.30 |
| Phy | Cortex Lab | Version 2.0b6 |
| GraphPad Prism | GraphPad Software | Version 10.6.0 |
| R | R Foundation for Statistical Computing | Version 4.6.1 |
| Python | Python Software Foundation | Version 3.13.14 |
| Custom HMM, Neuropixels, and statistical analysis code | This paper | Provided as a reviewer-accessible submission file; repository DOI to be added before publication |
| **Other** | | |
| ALZET osmotic pump | ALZET | Model 2002; Order No. 0000296 |
| 32-channel silicon probe | NeuroNexus Technologies | A1x32-6mm-50-177-A32 |
| SmartBox Pro data-acquisition system | NeuroNexus Technologies | SmartBox Pro™ |
| Neuropixels 1.0 probe, metal cap | imec | PRB_1_4_0480_1_C |
| Neuropixels 1.0 headstage | imec | HS_1000 |

| REAGENT or RESOURCE | SOURCE | IDENTIFIER |
|---|---|---|
| Neuropixels cable | imec | CBL_1000 |
| Neuropixels PXIe acquisition system | imec | PXIe_1000 |
| USB multifunction I/O device | National Instruments | NI USB-6212 |
| Laser displacement sensor head | KEYENCE | Model LK-H022 |
| Ag/AgCl pellet electrode | World Precision Instruments | Order code EP1 |
| Bovie cautery replacement tip, fine, low-temperature | Bovie, Aspen Surgical | Part No. H100 |
| ProJet 6000 HD stereolithography 3D printer | 3D Systems | Model ProJet 6000 HD |

2
