## Supplementary Movie 1 Legend for "A reactivated thalamocortical plasticity window promotes learning and is reshaped by experience"

Supplementary Movie 1 | Performance of the head-fixed Go/No-go whisker-detection task. Representative hit, miss, false-alarm, and correct-rejection trials are shown. Indicator lights mark Go, No-go, and lick events. Related to Fig. 1.
